# Nuclear envelope buffering of cytoskeletal force stabilizes chromosome interactions in *C. elegans* meiosis

**DOI:** 10.64898/2026.09.07.749962

**Authors:** Rachel Nenstiel, Chenshu Liu

## Abstract

Meiosis requires coordination between chromosome and nuclear envelope (NE) dynamics. Cytoskeletal forces transmitted through the NE via Linker of Nucleoskeleton and Cytoskeleton (LINC) complexes drive chromosome movement but can also compromise nuclear integrity if not properly balanced. Here, we identify the conserved inner nuclear membrane protein NEMP-1 as a key regulator that contributes to buffering of excessive force at the meiotic NE in *Caenorhabditis elegans*. NEMP-1 localizes to the NE throughout meiotic prophase and becomes enriched at the NE in nuclei with a weakened nuclear lamina. Simultaneous depletion of NEMP-1 and the lamin protein LMN-1 results in premature nuclear collapse and defects in chromosome synapsis and homolog pairing. These phenotypes depend on excessive dynein-mediated forces acting on a compromised NE. Furthermore, analysis of homolog pairing in the absence of synapsis revealed that NEMP-1 and LMN-1 contribute to the stabilization of synapsis-independent homolog interactions at the NE. Together, our findings reveal that excessive force at the NE can destabilize homolog interactions, and support a model in which NEMP-1 and LMN-1 cooperatively buffer cytoskeletal forces to maintain nuclear integrity while stabilizing chromosome interactions during meiosis.

**One-sentence summary:** Two nuclear envelope proteins synergize to maintain homolog interactions and meiotic nuclear integrity.

## Introduction

Meiosis is a specialized cell division program that partitions the diploid genome to make haploid gametes such as sperm and eggs (1). Many sexually reproducing organisms have an extended meiotic prophase during which homologous chromosomes must pair with each other and establish physical interactions in a timely manner (2). This process includes the initial pairing between homologs and the formation of the synaptonemal complex (SC) – a process known as synapsis – along the entire lengths of paired homologous chromosomes, both of which are prerequisite for recombination and the subsequent segregation of these chromosomes during meiotic metaphase (3–8).

Across the majority of eukaryotes, the meiotic nuclear envelope (NE) plays important roles in the establishment of homolog interactions (9–11). To facilitate homolog pairing, chromosomes often use specialized chromosomal regions, such as telomeres, to tether to the NE, where clustered proteins such as the linker of nucleoskeleton and cytoskeleton (LINC) complexes transduce cytoskeleton-mediated mechanical force to drive chromosomal movement (12–27). Rapid prophase movement (RPM) of chromosomes is a widely conserved phenomenon and is considered to facilitate the search for homologs and their subsequent pairing along the two-dimensional surface of the NE, promoting homolog interactions (28; 29).

*Caenorhabditis elegans* is a premier model organism for studying the spatiotemporal regulation of meiosis (7; 30; 31). In *C. elegans*, homolog pairing depends on specialized chromosomal regions near the end of each chromosome called pairing centers (PCs) (28; 32). PCs associate with LINC complex clusters at the NE, which transmit microtubule-based forces to drive chromosome movement, promoting pairing and synapsis initiation (25; 29; 33; 34). However, mechanical forces transmitted through the NE via LINC complexes can also compromise nuclear integrity if not properly balanced, leading to acute nuclear collapse in maturing oocytes (35; 36). Thus, the meiotic NE must both transmit and resist mechanical force. How meiotic nuclei buffer cytoskeletal forces while promoting chromosome movement, homolog interactions, and nuclear integrity remains poorly understood.

A clue to how this balance is achieved emerged from studies of the *C. elegans* synapsis checkpoint (37). When synapsis fails to complete, nuclear lamina, which in *C. elegans* is composed of the single lamin protein LMN-1(38), is greatly weakened, a step required for asynapsis-induced apoptosis to remove defective oocytes (37; 39). However, because apoptosis takes hours to complete (40; 41), whereas meiotic nuclei can collapse within minutes in the absence of lamin (35; 36), a paradox arises: how do meiotic nuclei maintain their integrity when synapsis checkpoint is activated? Here we identify the Nuclear Envelope Integral Membrane Protein 1 (NEMP-1) as a key regulator that helps resolve this paradox.

## Results and discussion

### NEMP-1 localizes to the meiotic nuclear envelope and intensifies when lamin is weakened

To understand how meiotic nuclei resist collapse when lamin is diminished during synapsis checkpoint activation, we hypothesized that additional NE proteins contribute to nuclear stability. We focused on the inner nuclear membrane protein NEMP-1, which is conserved in worms, flies, zebrafish, mice, and humans (42). Loss-of-function mutations of nemp-1 orthologs often result in reduced fertility across organisms, and human single-nucleotide polymorphisms of *NEMP1* gene are linked to premature menopause (42).

NEMP-1 is predicted to be an integral membrane protein with multiple membrane-spanning regions (Figure 1A). To characterize NEMP-1 function in *C. elegans* oogenesis, we epitope-tagged endogenous NEMP-1 and examined its localization using indirect immunofluorescence. NEMP-1 localizes to the meiotic NE throughout prophase (Figure 1, B and C). Strikingly, in prophase nuclei where synapsis fails to complete, either spontaneously in “straggler” nuclei (37; 43–45) or upon inhibiting the assembly of the synaptonemal complex (SC) by *syp-2(RNAi)* (37), NEMP-1 levels at the NE significantly increased concomitantly with weakened LMN-1 staining (Figure 1, D to F; Figure S1, A and B). The anticorrelation between NEMP-1 and LMN-1 levels at the NE is reminiscent of the complementary expression of Nemp1 and LaminA/C at the oocyte NE during mouse oogenesis (42). While the nature of this anticorrelation remains to be elucidated, these observations raised the possibility that increased NEMP-1 at the NE contributes to nuclear stability when lamin is diminished.

**Figure 1.**
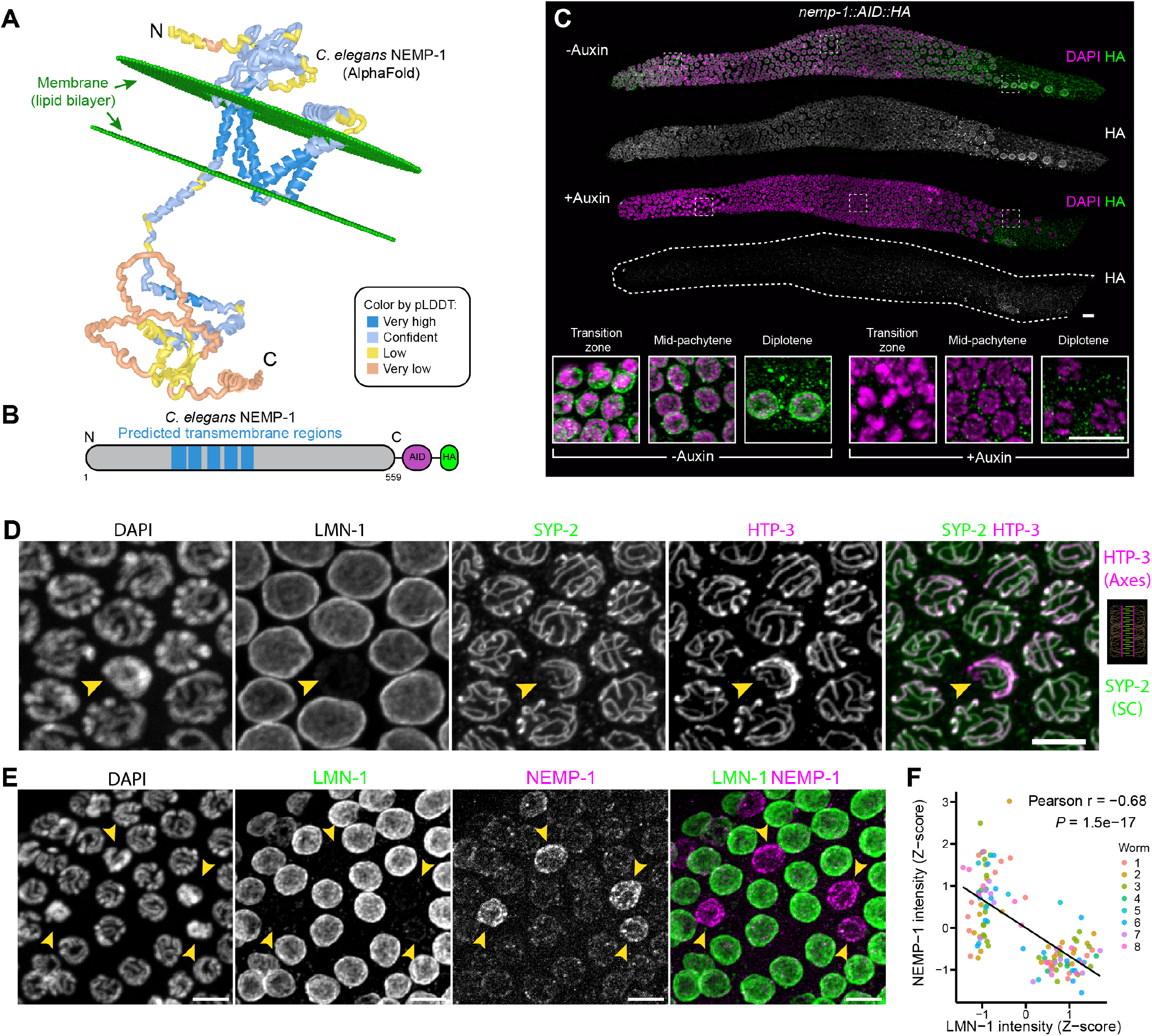
*C. elegans* NEMP-1 localizes to the meiotic nuclear envelope (NE) and is enriched at the NE of straggler nuclei with synapsis defects. **(A)** Predicted structure and membrane orientation of NEMP-1. AlphaFold-predicted structure of *C. elegans* NEMP-1 positioned within an ER-like membrane using the PPM 3.0 server. The structure is colored by AlphaFold per-residue confidence (pLDDT): dark blue, very high confidence (>90); light blue, confident (70–90); yellow, low confidence (50–70); and orange, very low confidence (<50). The lipid bilayer shows the predicted positioning of NEMP-1 within the membrane according to PPM 3.0. The membrane is depicted as a lipid bilayer, with the two green disks representing its opposing lipid headgroup layers; the bilayer is shown in perspective. **(B)** Schematic of *C. elegans* NEMP-1, indicating the predicted transmembrane regions and the C-terminal auxin-inducible degron (AID) and HA epitope tags. (C) Immunofluorescence images of NEMP-1::AID::HA in dissected gonads from adult hermaphrodites. DAPI is shown in magenta and HA in green or grayscale. Scale bars, 10 µm. (D) Immunofluorescence images of DAPI, lamin (LMN-1), chromosome axes (visualized by HTP-3) and synaptonemal complex (SC, visualized by SYP-2) staining in pachytene nuclei. SYP-2 in green and HTP-3 in magenta in the merged image. The yellow arrowhead indicates a “straggler” nucleus with incomplete synapsis and reduced LMN-1 staining. Scale bar, 5 µm. (E) Immunofluorescence images of DAPI, LMN-1 and NEMP-1 staining in pachytene nuclei. LMN-1 in green and NEMP-1 in magenta in the merged image. Yellow arrowheads indicate straggler nuclei. Scale bars, 5 µm. (F) Analysis of the relationship between LMN-1 and NEMP-1 intensities at the pachytene NE. LMN-1 and NEMP-1 intensities were Z-score normalized within each animal to account for inter-animal differences in fluorescence intensity and dynamic range. Normalized measurements of 122 nuclei pooled from eight animals were used for visualization and Pearson correlation analysis. LMN-1 and NEMP-1 intensities were also negatively correlated in each of the eight animals examined. Exact Pearson correlation coefficient and the P value are indicated in the figure.

### NEMP-1 and lamin cooperate to prevent premature prophase nuclear collapse

To test whether NEMP-1 is essential for stabilizing meiotic nuclei when LMN-1 is absent, we used auxininducible degradation (46; 47) to deplete NEMP-1 alone or together with LMN-1 in the germline and examine nuclear morphology (Figure 1B). Endogenous NEMP-1 was tagged with an auxin-inducible degron (AID) and an HA epitope at the C-terminus, and the effective loss of HA staining following auxin treatment is consistent with the C terminus of NEMP-1 facing the nucleoplasm, where it is accessible to germline-expressed TIR1 (Figure 1, B and C; Figure S1, C to E). The *C. elegans* germline provides a spatiotemporal gradient for analyzing how nuclear morphology changes as a function of time, as meiosis progresses from the mitosis-to-meiosis transition (“transition zone”) through pachytene stage where synapsis completes (Figure 1C; Figure 2, A and B). By dividing age-matched adult gonads into six equal-length zones and quantifying the percent of nuclei with hypercondensed DAPI staining in each zone, we observed a striking synthetic effect of depleting NEMP-1 and LMN-1: while depletion of NEMP-1 or LMN-1 alone did not overtly affect pachytene nuclear morphology, simultaneous depletion of NEMP-1 and LMN-1 caused a dramatic increase in the fraction of pachytene nuclei with hypercondensed chromatin (Figure 2, A and B). To determine whether the reduced nuclear size reflected acute collapse rather than impaired nuclear growth, we performed time-lapse live imaging using the LINC complex component ZYG-12::GFP as a marker (Figure 2C; and Video 1). Indeed, nuclear contours visualized by ZYG-12::GFP revealed abrupt nuclear collapse occurring within approximately one minute, followed by stabilization at a substantially reduced nuclear size (Figure 2, D and E), supporting that NEMP-1 and LMN-1 work together to prevent premature nuclear collapse in pachytene.

**Figure 2.**
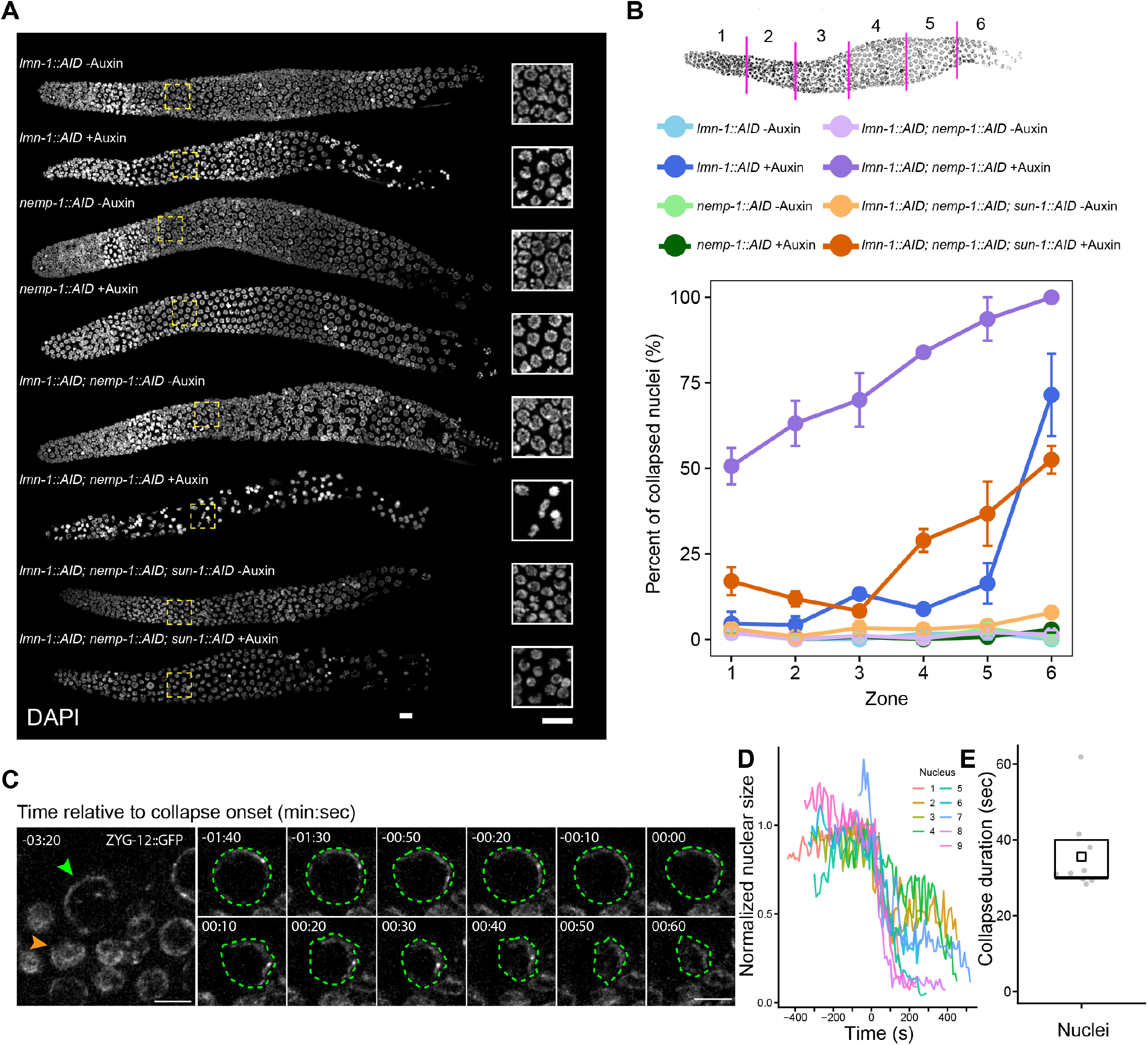
NEMP-1 and LMN-1 cooperate to prevent premature prophase nuclear collapse. (A) Nuclear morphology visualized by DAPI staining throughout meiotic prophase. All animals were homozygous for *Psun-1::TIR1* or *Pgld-1::TIR1* (omitted in the figure). Insets show early pachytene nuclei. Scale bars, 10 µm. (B) Percent of collapsed nuclei as a function of meiotic progression. Schematics at the top show distal gonad being divided into six zones of equal lengths, so that the percent of nuclei with collapsed/condensed chromatin in each zone can be quantified. Three animals were analyzed per condition, and data are plotted as mean ± SEM. Nuclear counts were pooled across animals for pairwise comparisons of proportions. Comparisons are made for dark purple vs. light purple, dark green, dark blue, and dark orange, respectively: zone 1: P = 8.30e-27, 6.48e-28, 1.83e-23, and 8.60e-09; zone 2: P = 1.90e-42, 1.86e-43, 1.26e-27, and 2.26e-15; zone 3: P = 1.32e-40, 2.27e-43, 5.56e-21, and 6.57e-24; zone 4: P = 3.80e-59, 2.94e-63, 2.51e-40, and 6.56e-18; zone 5: P = 4.27e-60, 9.79e-70, 7.97e-32, and 1.47e-17; zone 6: P = 1.72e-43, 2.20e-59, 1.82e-07, and 8.77e-13. (C) Representative time-lapse images showing a collapsing pachytene nucleus after depletion of NEMP-1 and LMN-1. Maximum-intensity projections were scaled identically. Time stamps are min:sec. The green arrowhead indicates the pachytene nucleus before collapse, and the orange arrowhead indicates a collapsed nucleus. The green dashed line outlines the collapsing nucleus. Scale bar, 5 µm. (D) Quantification of nuclear size change during nuclear collapse measured from timelapse ZYG-12::GFP images. Nine collapsing nuclei from eight animals were measured. Nuclear collapse kinetics were quantified from time-lapse images acquired at 10-s intervals. Nuclear size was normalized to the pre-collapse size, and trajectories were aligned such that t = 0 corresponded to the measured time point at which normalized nuclear size was closest to 1. (E) Collapse duration was measured for the nuclei analyzed in D from sustained size reduction to subsequent stabilization. Points represent individual nuclei; the box indicates the median and interquartile range, and the open square indicates the mean.

In *C. elegans*, constitutive *nemp-1* deletion mutants have markedly reduced fertility, especially at 15°C (42). Continuous AID-mediated depletion of NEMP-1 in the germline, with auxin treatment beginning at the L1 stage, did not measurably reduce cumulative brood size at either 20°C or 15°C, although egg laying was delayed and embryonic viability was modestly reduced, with a stronger effect at 15°C (Table S1, Figure S2A). Thus, while germline-restricted depletion of NEMP-1 did not fully recapitulate the pronounced reproductive defects previously reported for constitutive *nemp-1* deletion alleles (42), this difference may reflect aspects of NEMP-1 function that are not fully captured by our AID-based depletion strategy.

### NEMP-1 and lamin cooperate to stabilize interactions between homologous chromosomes

While depletion of NEMP-1 or LMN-1 alone did not overtly affect homolog pairing or synapsis (Figure 3, A to D), simultaneous depletion of NEMP-1 and LMN-1 resulted in distinct phenotypes in homolog pairing and synapsis. When scoring the percent of nuclei with paired X-chromosome PCs, marked by the zinc finger PC protein HIM-8, as a function of meiotic progression, we found that in animals depleted of both NEMP-1 and LMN-1, homologs largely achieved pairing before subsequently separating (Figure 3, A and B; Figure S2B), indicating that NEMP-1 and LMN-1 are particularly important for the stable maintenance of homolog pairing over meiotic progression. Notably, when quantifying the percent of nuclei with complete synapsis as a function of meiotic progression (Figure 3, C and D; Figure S2C), we found that nuclei exhibiting transient homolog pairing already showed severe synapsis defects, indicating that either synapsis defects precede pairing instability, or that synapsis is particularly sensitive to simultaneous loss of NEMP-1 and LMN-1.

**Figure 3.**
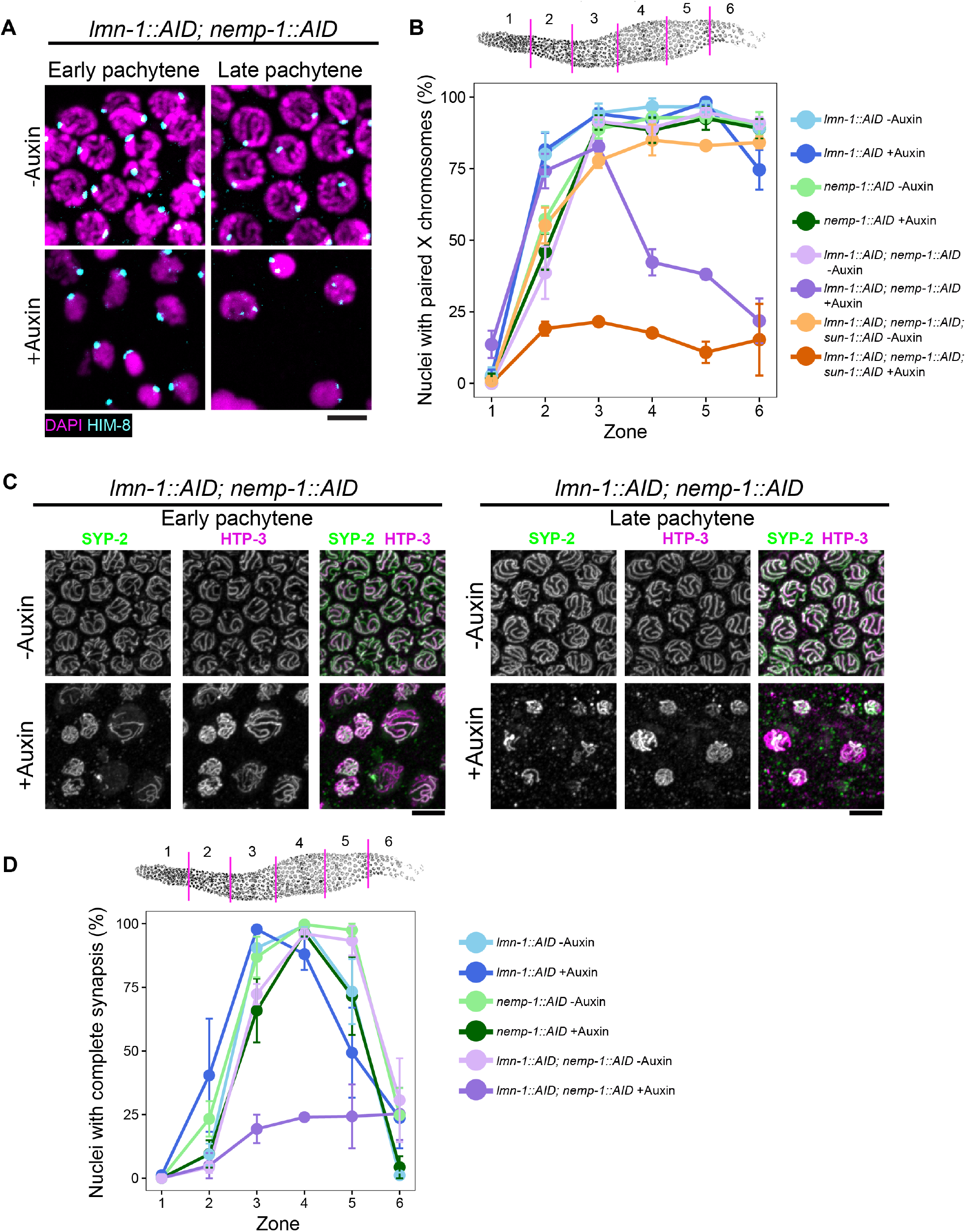
NEMP-1 and lamin cooperate to stabilize interactions between homologous chromosomes. (A) Immunofluorescence images showing X-chromosome homolog pairing visualized by HIM-8 staining. HIM-8 in cyan and DAPI in magenta. Scale bar, 5 µm. (B) Quantification of the percent of nuclei with paired X-chromosome pairing centers as a function of meiotic progression. Distal gonad was divided into six equal-length zones, and the percent of nuclei with a single HIM-8 focus, indicating paired X-chromosome PCs, in each zone was quantified. Three animals were analyzed per condition, and data are plotted as mean ± SEM. Nuclear counts were pooled across animals for pairwise comparisons of proportions. Comparisons are made for dark purple vs. light purple, dark green, dark blue, and dark orange, respectively: zone 3: P = 1.74e-02, 3.63e-02, 2.74e-03, and 2.43e-22; zone 4: P = 6.07e-18, 2.54e-16, 3.58e-20, and 2.16e-05; zone 5: P = 6.10e-25, 6.37e-23, 4.15e-25, and 1.09e-05; zone 6: P = 1.93e-19, 1.64e-25, 6.00e-10, and 5.41e-01. Comparisons are also made for dark orange vs. light orange: zone 3: P= 3.80e-28; zone 4: 5.48e-41; zone 5: 5.37e-36; zone 6: 1.37e-19. (C) Immunofluorescence images showing chromosome synapsis visualized by SYP-2 and HTP-3 staining. SYP-2 in green and HTP-3 in magenta. Incomplete synapsis was identified as regions of chromosomes that have HTP-3 but lack SYP-2 staining. Scale bar, 5 µm. (D) Quantification of the percent of nuclei with complete synapsis as a function of meiotic progression. Distal gonad was divided into six equal-length zones, and the percent of nuclei with complete synapsis (defined as all HTP-3-marked chromosome axes having colocalized SYP-2 staining) in each zone was quantified. Three animals were analyzed per condition, and data are plotted as mean ± SEM. Nuclear counts were pooled across animals for pairwise comparisons of proportions. Comparisons are made for dark purple vs. light purple, dark green, and dark blue, respectively: zone 3: P = 1.95e-15, 8.24e-13, and 2.75e-37; zone 4: P = 3.11e-36, 2.09e-41, and 1.00e-17; zone 5: P = 7.90e-23, 5.04e-11, and 1.52e-02; zone 6: P = 8.34e-01, 1.61e-03, and 1.00e+00.

### Simultaneous loss of NEMP-1 and lamin increases LINC complex dynamics at the NE

Because homolog pairing and synapsis are promoted by chromosome movements driven through PC-associated LINC complexes, we next asked whether NEMP-1 and LMN-1 influence LINC complex dynamics at the meiotic NE. LINC complexes (shown by ZYG-12::GFP) normally exhibit prominent clustering in the transition zone, concomitant with rapid chromosome movement during homolog pairing and synapsis initiation (24; 25; 29; 36; 48; 49). While depletion of NEMP-1 alone did not change the duration of LINC clustering, simultaneous depletion of NEMP-1 and LMN-1 substantially prolonged LINC complex clustering at the NE, consistent with prolonged engagement of the chromosome-movement program (Figure 4, A to C). NEMP-1 and LMN-1 co-depletion also prolongs the region of meiotic prophase where nuclei stain positive for pHIM/ZIM, a readout of CHK-2 activity (43), consistent with prolonged CHK-2 activity and delayed exit from the chromosome-movement-competent state (Figure S2, D and E).

**Figure 4.**
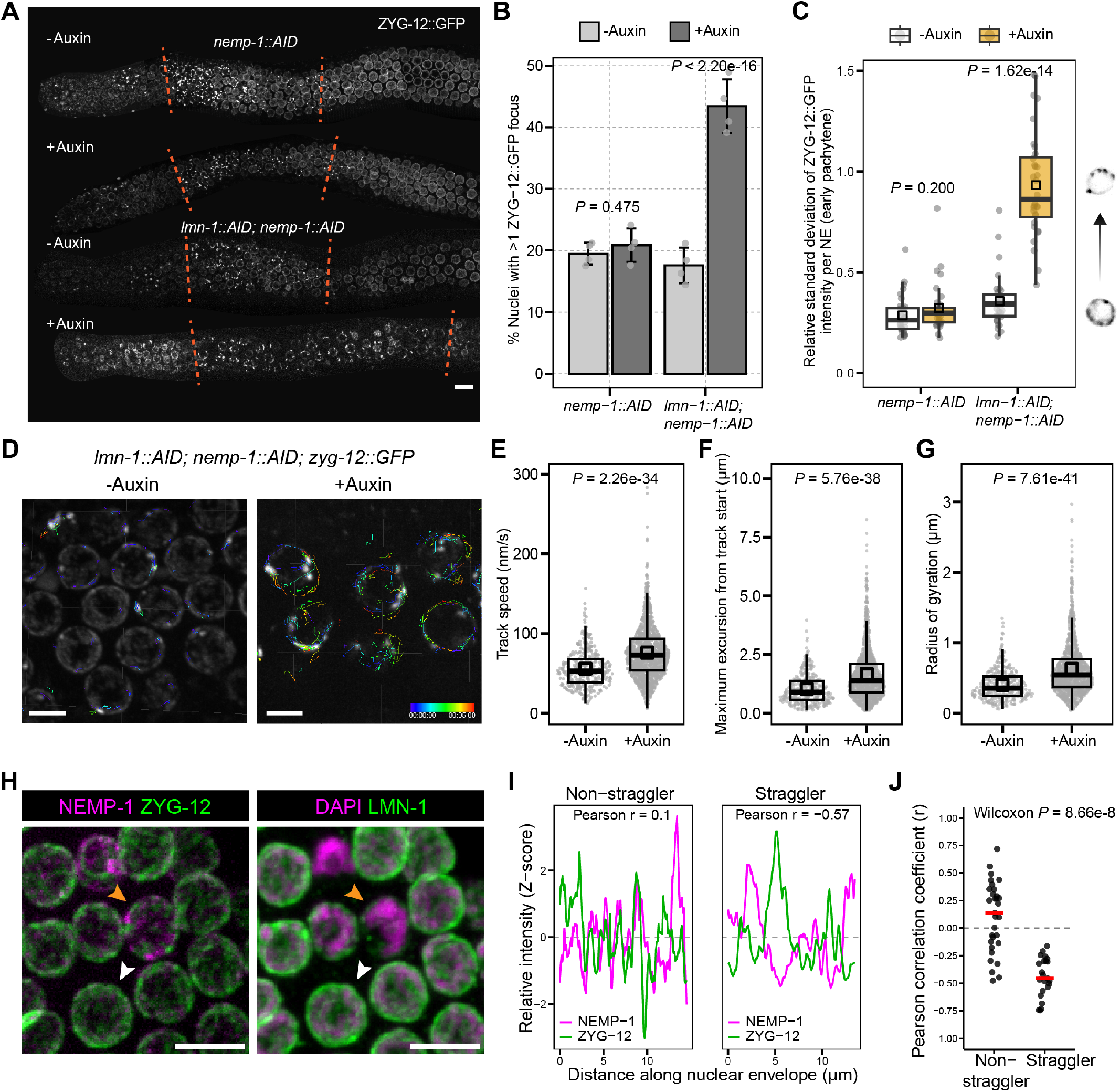
Simultaneous loss of NEMP-1 and LMN-1 increases LINC complex dynamics at the NE. (A) Grayscale maximum-intensity Z-projection images from 3D live imaging showing prolonged ZYG-12::GFP clustering at the NE following simultaneous depletion of NEMP-1 and LMN-1. Red dashed lines indicate the region of the gonad in which meiotic nuclei exhibit ZYG-12::GFP clustering at the NE. Scale bar, 10 µm. (B) Quantification of the percent of prophase nuclei with more than one ZYG-12::GFP focus at the NE. Four animals were analyzed per condition. Nuclear counts were pooled across animals for pairwise comparison of proportions. Exact P values are indicated in the figure. (C) Clustering of ZYG-12::GFP, defined as the relative standard deviation (ratio of the standard deviation to the mean value, also known as coefficient of variation) of fluorescence intensity at the NE in each nucleus. Medians (black crossbars) and means (black boxes) are shown. Fluorescence intensity surrounding each nucleus was manually segmented and quantified from additive projection images after background subtraction. For each genotype, 32 nuclei were pooled from 4 animals for the control group, and 32 nuclei were pooled from 4 animals for the auxin treatment group. Unpaired Welch t-test was used to compute the P values. Exact P values are indicated in the figure. (D) Grayscale images of ZYG-12::GFP overlaid with color-coded trajectories generated by 3D particle tracking of ZYG-12::GFP patches. Scale bar, 4 µm. (E - G) Dynamics of ZYG-12::GFP patches at the early pachytene NE, quantified as the mean speed of individual patches (E), maximum excursion distance from each track’s origin (F), and the radius of gyration of each track (G). All measurements were calculated after drift correction. Medians (black crossbars) and means (black boxes) are shown. 335 clusters of ZYG-12::GFP were pooled from 4 animals for the control group, and 1287 clusters were pooled from four animals for the auxin treatment group. P values were calculated with Welch two sample t-test and are included in the figure. (H) Immunofluorescence images (maximum-intensity partial Z-projections) showing the spatial relationship between NEMP-1 and ZYG-12 at the NE in pachytene nuclei. The orange arrowhead indicates a straggler nucleus, and the white arrowhead indicates a non-straggler nucleus. Scale bar, 5 µm. (I) Representative line scans (after Z-score normalization) showing how NEMP-1 intensities covary with ZYG-12 intensities as a function of distance along the meiotic NE. Additive partial Z-projection images were used for line scan analysis. (J) Quantification of the Pearson correlation coefficients between NEMP-1 and ZYG-12 in non-stragglers and stragglers. 31 non-straggler nuclei from six animals and 23 straggler nuclei from eight animals were analyzed. Red bars indicate the medians. A Wilcoxon rank-sum test with continuity correction was used to compute the P value. The exact P value is indicated in the figure.

Importantly, using live imaging and three-dimensional particle tracking of ZYG-12 clusters over time in early pachytene nuclei, we found significantly increased cluster motility following co-depletion of NEMP-1 and LMN-1, suggesting that these NE components normally constrain LINC complex dynamics and, consequently, the spatial transmission of cytoskeletal forces across the NE (Figure 4, D and E; and Video 2). Indeed, following NEMP-1/LMN-1 co-depletion, individual ZYG-12 clusters exhibited increased maximum excursion from their starting positions and increased radius of gyration, indicating that they explored a larger spatial domain of the NE (Figure 4, F and G). Consistent with this possibility, in spontaneously occurring straggler nuclei, where polarized chromosomes and weakened LMN-1 often indicate synapsis checkpoint activation (Figure 1D) (37), NEMP-1 and ZYG-12 occupied largely non-overlapping domains of the NE (Figure 4, H to J). Together, the increased spatial exploration of ZYG-12 clusters following NEMP-1/LMN-1 depletion and the spatial segregation of NEMP-1 and ZYG-12 in straggler nuclei raise the possibility that NEMP-1 contributes to spatial confinement of LINC complexes at the meiotic NE.

### Excessive mechanical forces at the meiotic nuclear envelope destabilize nuclear integrity and homolog interactions

During meiotic prophase, LINC complexes comprising of the SUN-1/ZYG-12 bridge (SUN-1 at the inner nuclear membrane and ZYG-12 at the outer nuclear membrane) engage cytoplasmic dynein to transmit microtubule-dependent forces across the NE (29; 36). This force-transmission machinery is most active during the transition zone, where it drives chromosome movement, and normally subsides as homologs pair and synapse (29; 48). When synapsis defects persist, however, LINC complexes and associated proteins remain clustered at the NE for an extended period, prolonging the chromosome-movement-competent state and continued transmission of cytoskeletal forces (25; 29; 37; 50–52). Under these conditions, the concomitant reduction in LMN-1 weakens the mechanical properties of the meiotic NE (36; 37), potentially increasing its reliance on other NE components to withstand continued cytoskeletal force. We therefore asked whether, in the additional absence of NEMP-1, LINC-mediated forces become excessive relative to the mechanical buffering capacity of the NE, thereby driving premature nuclear collapse and destabilization of homolog interactions. To test this possibility, we first perturbed mechanotransduction through the LINC complex by induced degradation of SUN-1. Indeed, simultaneous depletion of SUN-1 significantly suppressed premature nuclear collapse caused by co-depletion of NEMP-1 and LMN-1, supporting the conclusion that LINC-transmitted mechanical forces drive premature collapse of mechanically compromised nuclei (Figure 2, A and B).

However, because SUN-1 depletion also causes non-homologous synapsis and pairing defects (Figure 3B) (24; 25; 36; 49), we used RNAi to deplete DNC-1 (dynactin), an activator of cytoplasmic dynein (53–57). Cytoplasmic dynein is known to be important for mechanotransduction across meiotic NE but whose depletion does not itself cause major homolog-pairing defects (25; 29; 36; 47). Indeed, simultaneous depletion of DNC-1 not only suppressed premature nuclear collapse caused by co-depletion of NEMP-1 and LMN-1 (Figure 5, A and B), but also largely restored sustained homolog pairing and synapsis even when both NEMP-1 and LMN-1 are absent (Figure 5, C to F). These data suggest that dynein-mediated forces acting on a mechanically compromised NE contribute to the destabilization of both nuclear integrity and homolog interactions.

**Figure 5.**
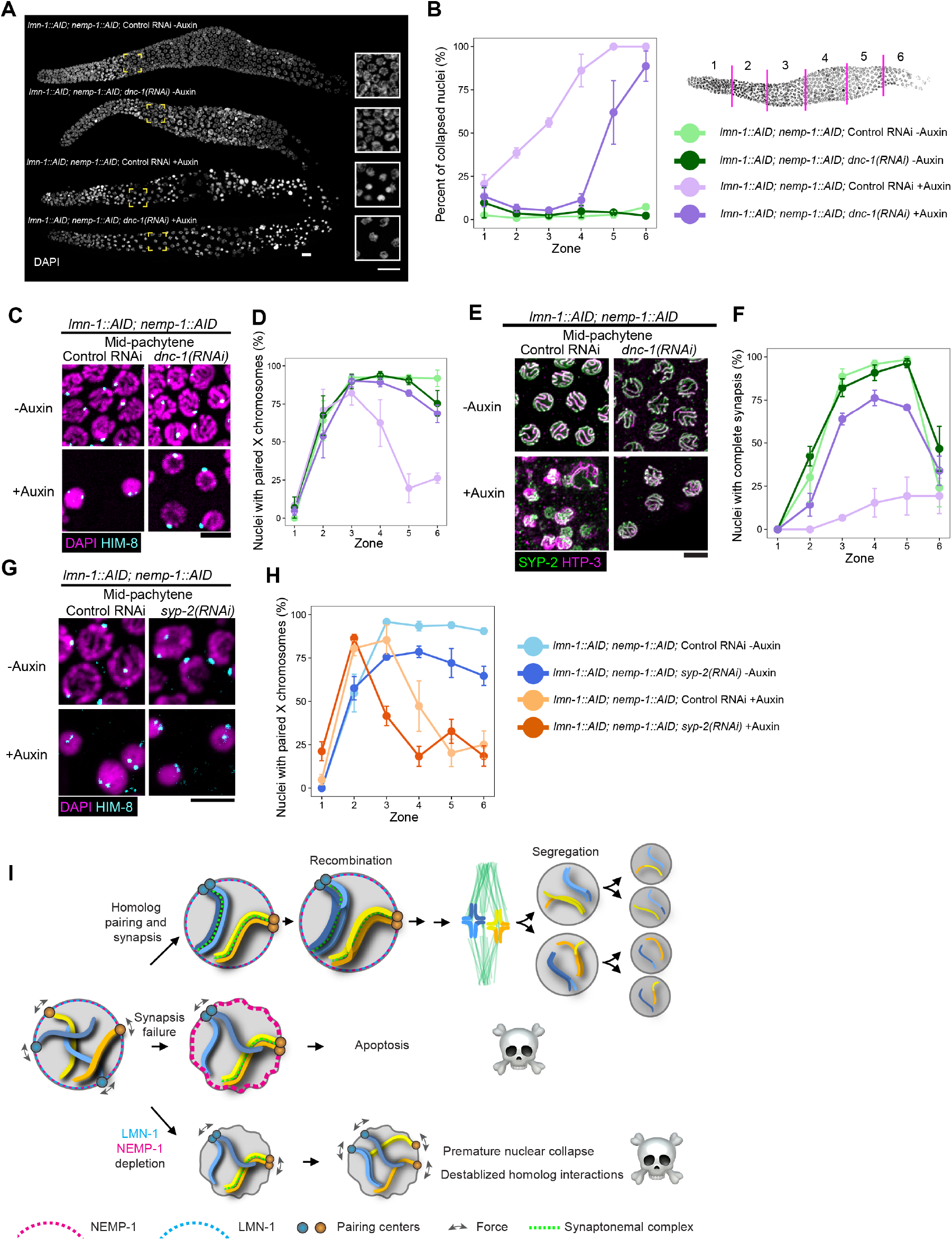
Excessive dynein-mediated forces at the meiotic nuclear envelope destabilize nuclear integrity and homolog interactions. (A) Nuclear morphology visualized by DAPI staining throughout meiotic prophase. All animals were homozygous for *Psun-1::TIR1* or *Pgld-1::TIR1* (omitted in the figure). Insets show early pachytene nuclei. Scale bars, 10 µm. (B) Percent of collapsed nuclei as a function of meiotic progression. Schematics at the top right shows distal gonad being divided into six zones of equal lengths, so that the percent of nuclei with collapsed/condensed chromatin in each zone can be quantified. Three animals were analyzed per condition, and data are plotted as mean ± SEM. Nuclear counts were pooled across animals for pairwise comparisons of proportions. Comparisons are made for dark purple vs. light purple and dark purple vs. dark green, respectively: zone 2: P = 8.82e-11 and 1.96e-01; zone 3: P = 1.04e-18 and 2.35e-01; zone 4: P = 6.91e-34 and 2.34e-02; zone 5: P = 1.57e-15 and 1.66e-33; zone 6: P = 2.11e-03 and 6.22e-48. (C) Immunofluorescence images showing X-chromosome homolog pairing visualized by HIM-8 staining. HIM-8 in cyan and DAPI in magenta. Scale bar, 5 µm. (D) Quantification of the percent of nuclei with paired X-chromosome pairing centers as a function of meiotic progression. Distal gonad was divided into six equal-length zones (as illustrated in B), and the percent of nuclei with a single HIM-8 focus, indicating paired X-chromosome PCs, in each zone was quantified. Three animals were analyzed per condition, and data are plotted as mean ± SEM. Nuclear counts were pooled across animals for pairwise comparisons of proportions. Comparisons are made for dark purple vs. light purple, dark purple vs. dark green, and light green vs. light purple, respectively: zone 3: P = 0.12, 0.73, and 0.01; zone 4: P = 1.15e-07, 0.08, and 2.29e-13; zone 5: P = 1.77e-23, 0.04, and 2.53e-43; zone 6: P = 1.53e-10, 0.07, and 9.29e-23. Color-coding of genotypes and experimental conditions are identical to those in B. (E) Immunofluorescence images showing chromosome synapsis visualized by SYP-2 and HTP-3 staining. SYP-2 in green and HTP-3 in magenta. Incomplete synapsis was identified as regions of chromosomes that have HTP-3 but lack SYP-2 staining. Scale bar, 5 µm. (F) Quantification of the percent of nuclei with complete synapsis as a function of meiotic progression. Distal gonad was divided into six equal-length zones (as illustrated in B), and the percent of nuclei with complete synapsis (defined as all HTP-3-marked chromosome axes having colocalized SYP-2 staining) in each zone was quantified. Three animals were analyzed per condition, and data are plotted as mean ± SEM. Nuclear counts were pooled across animals for pairwise comparisons of proportions. Comparisons are made for dark purple vs. light purple, dark purple vs. dark green, and light green vs. light purple, respectively: zone 2: P = 2.09e-05, 1.20e-07, and 1.09e-13; zone 3: P = 1.37e-16, 1.35e-04, and 7.31e-43; zone 4: P = 1.61e-18, 3.99e-04, and 1.42e-48; zone 5: P = 2.31e-08, 7.38e-10, and 1.83e-37; zone 6: P = 0.18, 0.08, and 0.55. Color-coding of genotypes and experimental conditions are identical to those in B. (G) Immunofluorescence images showing X-chromosome homolog pairing visualized by HIM-8 staining. HIM-8 in cyan and DAPI in magenta. Scale bar, 5 µm. (H) Quantification of the percent of nuclei with paired X-chromosome pairing centers as a function of meiotic progression. Distal gonad was divided into six equal-length zones (as illustrated in B), and the percent of nuclei with a single HIM-8 focus, indicating paired X-chromosome PCs, in each zone was quantified. Three animals were analyzed per condition, and data are plotted as mean ± SEM. Nuclear counts were pooled across animals for pairwise comparisons of proportions. Comparisons are made for dark orange vs. light orange and dark orange vs. dark blue, respectively: zone 2: P = 2.83e-01 and 4.00e-08; zone 3: P = 1.83e-11 and 1.16e-10; zone 4: P = 1.41e-04 and 1.18e-21; zone 5: P = 1.92e-01 and 4.58e-11; zone 6: P = 3.21e-01 and 2.43e-13. Comparisons are also made for light blue vs. dark blue. Zone 3: P = 7.73e-13; zone 4: P = 6.65e-09; zone 5: P = 1.11e-11; zone 6: P = 7.62e-09. (I) Model for cooperative buffering of cytoskeletal force by NEMP-1 and LMN-1 to maintain homolog interactions and meiotic nuclear integrity. When LMN-1 is diminished following defective synapsis, NEMP-1 becomes enriched at the NE. Simultaneous loss of LMN-1 and NEMP-1 compromises meiotic nuclear integrity and homolog interactions through excessive dynein-mediated force.

Finally, to understand how excessive forces lead to destabilized homolog interactions, in particular, whether pairing defects are secondary to synapsis defects in the absence of NEMP-1 and LMN-1, we analyzed homolog pairing in the absence of synapsis using *syp-2(RNAi)* (Figure 5, G and H). Indeed, we observed that when both NEMP-1 and LMN-1 are present, the absence of synapsis itself did not cause the transient pairing phenotype but led to a plateau at approximately 70% nuclei with paired HIM-8, consistent with previous findings of synapsis-independent pairing (58–62). Notably, when synapsis is defective [*syp-2(RNAi)*], simultaneous depletion of NEMP-1 and LMN-1 still produced transient pairing followed by rapid loss of pairing maintenance (Figure 5, G and H). These results suggest that NEMP-1 and LMN-1 promote synapsis-independent maintenance of homolog pairing at the NE. Our data also suggest that synapsis contributes to homolog-pairing stability even in the absence of NEMP-1 and LMN-1. Mammalian Nemp1 was recently shown to be important for synapsis (63). How *C. elegans* NEMP-1 and LMN-1 stabilize homolog pairing and synapsis against excessive force remains to be elucidated.

### Nuclear envelope–mediated buffering of cytoskeletal forces stabilizes homolog interactions

Together, our findings illustrate how NEMP-1 and LMN-1 cooperate to balance the need for the meiotic NE to both transmit and resist force during the extended prophase (Figure 5I). During normal meiosis, NEMP-1 and LMN-1 coexist at the NE during homolog pairing and synapsis. When synapsis fails, LMN-1 is diminished as part of the synapsis checkpoint response, whereas NEMP-1 becomes enriched at the NE. We propose that this reciprocal response helps preserve nuclear integrity during the prolonged interval preceding checkpoint-induced apoptosis. However, when both NEMP-1 and LMN-1 are depleted, the meiotic NE becomes mechanically compromised, resulting in premature pachytene nuclear collapse and destabilization of homolog interactions. Both phenotypes are suppressed by reducing dynein-mediated force, supporting a model in which NEMP-1 and LMN-1 cooperatively buffer cytoskeletal forces at the meiotic NE.

While the relationship between nuclear collapse and homolog interaction defects remains unresolved, our finding that reducing dynein-mediated force transmission suppresses both phenotypes indicates that both depend on mechanical force. Persistent synapsis defects are known to prolong LINC complex clustering and chromosome movement (25; 29; 51; 52); however, the substantial restoration of synapsis following DNC-1 depletion places the synapsis defects caused by simultaneous loss of NEMP-1 and LMN-1 at least in large part down-stream of dysregulated mechanical force at the meiotic NE.

More broadly, the opposing changes in LMN-1 and NEMP-1 following synapsis failure suggest that the composition and mechanical state of the NE are dynamically remodeled in response to meiotic checkpoint activation (37). Given that human NEMP1 contributes to NE mechanical stiffness (42), NEMP-1 enrichment may provide compensatory mechanical support when LMN-1 is reduced, allowing nuclei to withstand prolonged chromosome-movement-associated forces.

Our work extends the established role of cytoskeletal force in promoting homolog interactions through RPM (64–67) by revealing a complementary requirement for the NE to withstand this mechanical load: when insufficiently buffered, the same forces that promote chromosome movement can instead destabilize homolog interactions. Collectively, our findings identify NEMP-1 and LMN-1 as components of this mechanical buffering system, conferring NE resistance to dynein-mediated force. These findings support a role for the meiotic NE as an active mechanical regulator, helping determine how cytoskeletal forces influence chromosome behavior while preserving nuclear integrity throughout meiotic prophase.

## Materials and Methods

### Generation of worm strains

All *C. elegans* strains were maintained at 20°C under standard growth conditions (68). Alleles generated in this study are detailed in Table S2, and a complete list of strains used and generated is provided in Table S3. Genome editing was performed using CRISPR-Cas9 with Alt-R CRISPR-Cas9 guide RNA (gRNA) products (Integrated DNA Technologies [IDT]), as previously described (36). Two independent *nemp-1::AID::HA* alleles, *emj1* and *emj2*, containing the same engineered sequence were generated and sequence-verified. The sequences of guide RNA and repair template are provided in Table S4.

### Fertility and viability

Brood size, progeny viability, and the frequency of males among the self-progeny of individual hermaphrodites were quantified as previously described (36). Briefly, L4 hermaphrodites (P0) were individually transferred onto 60-mm nematode growth medium (NGM) plates containing freshly spread, thin lawns of OP50. Animals were maintained at either 15°C or 20°C and transferred to fresh plates every 12 or 24 h for 5–6 days, or until egg laying ceased. To quantify brood size, viability, and male self-progeny following auxin treatment, age-matched hermaphrodites were maintained on auxin-containing plates beginning at the L1 stage following hatching from embryos. At the L4 stage, individual hermaphrodites were transferred to fresh auxin-containing plates and maintained using the same procedures described above.

### Auxin-inducible degradation (AID) and RNA interference (RNAi)

Plates containing 2 mM indole-3-acetic acid (IAA or auxin, I2886, Sigma) were prepared as previously described for the degradation of AID-tagged proteins (36). Unless otherwise indicated, young adult animals were exposed to auxin for 24 h prior to analysis. This duration was selected based on preliminary time-course experiments and was used consistently across single and combinatorial AID experiments (Figure S2; Figure 1–Figure 5). For live imaging experiments, animals were exposed to auxin for 12 h to capture nuclear collapse in real time or to assess LINC complex dynamics before the onset of widespread nuclear collapse. All animals used in AID experiments were homozygous for a germline-expressed TIR1 transgene driven by either the *sun-1* or *gld-1* promoter, as specified in Table S3; TIR1 transgenes are omitted from genotype labels in the figures for simplicity.

RNAi experiments were performed according to established protocols with minor modifications (36; 69; 70), using frozen stocks of HT115 bacteria carrying constructs from the Ahringer *C. elegans* RNAi feeding library (71; 72). For RNAi experiments, L4 hermaphrodites were transferred onto RNAi plates and dissected 48 h later. For experiments combining RNAi with auxin-inducible degradation, L4 hermaphrodites were placed on RNAi plates containing only IPTG first for 24 h and transferred again onto RNAi plates containing both IPTG and auxin to be dissected 24 h later. RNAi clone information is provided in Table S5. RNAi plasmid constructs were sequence verified with Plasmidsaurus Whole Plasmid Sequencing (Oxford Nanopore, R10.4.1).

### Immunofluorescence

Immunofluorescence was performed as previously described (31; 36). The following primary antibodies were used: anti-V5 (1:400; rabbit; V8137, Millipore-Sigma), anti-V5 (1:400; mouse; R96025, Invitrogen), anti-hemagglutinin (HA) (1:400; mouse monoclonal; 26183, Invitrogen), anti-SYP-2 (1:1000; affinity-purified rabbit) (58; 73), anti-pHIM/ZIM (1:1000; affinity-purified rabbit) (43), anti-HTP-3 (1:400; chicken) (61), and anti-HIM-8 (1:400; guinea pig) (28). Secondary antibodies conjugated to Alexa Fluor 488, 555, 568, and 647 were used at a 1:400 dilution (Jackson ImmunoResearch or Life Technologies). For fixed imaging, 3D image stacks were acquired using a spinning disk confocal (Yokogawa CSU-W1 standard disk paired with Zeiss Axio Observer 7 Basic MarianasTM Microscope, Intelligent Imaging Innovations (3i), Inc., Denver, CO, USA), with a 100x 1.46 NA oil immersion objective and a Prime 95B Back Illuminated Scientific CMOS camera at 1×1 binning, under the control of the SlideBook software (version 2025, 3i). 40 optical sections in Z with 0.2 µm spacing were acquired for each field of view. Acquisition parameters were held constant between matched control and experimental samples.

### Live imaging

Live imaging experiments were performed as previously described (36). To analyze LINC complex dynamics, 4D image stacks of gonadal region corresponding to early meiotic prophase were acquired every 5 s for a total of 60 time points. At each time point, 10 z-sections were acquired at 0.5-µm intervals. To examine the dynamics of nuclear collapse, 3D image stacks of the early pachytene region were acquired every 10 s for a total of 60 time points. Live imaging was performed on the same Marianas spinning-disk confocal system described above.

### Image Analysis

Movement of LINC complex clusters in 4D images was analyzed using Imaris (×64, version 9; Bitplane), as previously described (36). Maximum excursion was calculated as the greatest three-dimensional displacement from the initial track position, whereas radius of gyration was calculated as the root-mean-square three-dimensional distance of all track positions from the track centroid.

Analysis of nuclear envelope shrinking dynamics during nuclear collapse was performed using Fiji, as previously described (36). Nuclear collapse kinetics were quantified from time-lapse images acquired at 10-s intervals. Nuclear size measurements were normalized to the pre-collapse nuclear size for each nucleus. To compare trajectories among nuclei, the time point at which normalized nuclear size was closest to 1 was defined as t = 0, and all other time points were expressed relative to this reference point. Collapse duration was determined independently for each trajectory using a standardized algorithm designed to accommodate small frame-to-frame fluctuations: collapse onset was defined as the beginning of a sustained net decrease in normalized nuclear size, identified by a decrease of ≥ 0.03 over a two-frame (20-s) window. Following collapse onset, the end of collapse was defined as the first point at which nuclear size stabilized, operationally defined as a net decrease of <0.02 over a subsequent two-frame window. Collapse duration was calculated as the elapsed time between these onset and stabilization.

For fixed image analysis, 3D images were exported as 16-bit TIFF image stacks and projected and analyzed using Fiji. Homolog pairing, complete synapsis, and nuclear collapse across meiotic progression were quantified as previously described (36). Briefly, images spanning the distal premeiotic region through early diplotene were divided into six zones of equal length. Nuclei with one HIM-8 focus were scored as paired, whereas nuclei with two resolvable HIM-8 foci were scored as unpaired. Complete synapsis was scored when SYP-2 staining extended along all HTP-3-marked chromosome axes. Nuclear collapse in fixed images was quantified across the same six zones, and collapsed nuclei were identified by markedly reduced nuclear size and hypercondensed DAPI morphology, as previously described (36).

Maximum-intensity Z-projections of 3D image stacks were generated using SlideBook or Fiji for representative images in all figures. For all intensity-based comparisons, matched control and experimental images were displayed using identical lookup table (LUT) scaling. Quantitative intensity measurements of NEMP-1 (HA), LMN-1 (V5) or ZYG-12::GFP were performed on additive projection images after background subtraction as previously described (36), using manual segmentation and intensity measurements in Fiji/ImageJ (National Institutes of Health) or automated macro-based pipelines adapted from previously described algorithms (74). Quantitative analysis of ZYG-12::GFP clustering using relative standard deviation (also known as coefficient of variation) of intensities (Figure 4C) was performed as previously described (36).

### Modeling of NEMP-1 membrane positioning using AlphaFold and PPM3.0

The predicted structure of *C. elegans* NEMP-1 was obtained from the AlphaFold Protein Structure Database (UniProt Q19293; AlphaFold model version 6). To estimate the orientation of the predicted structure within a lipid bilayer, the AlphaFold model was submitted as a PDB file to the Positioning of Proteins in Membranes (PPM) 3.0 server (https://opm.phar.umich.edu/ppm_server3_cgopm) (75). The mammalian endoplasmic reticulum (ER) membrane model was used as the membrane environment, consistent with the continuity of the NE and ER. As a robustness check, positioning was also performed using the fungal ER membrane model and yielded a comparable predicted orientation. The PPM-positioned structure was visualized using iCn3D and colored according to AlphaFold predicted Local Distance Difference Test (pLDDT) scores, which report residue-level confidence in the predicted structure: >90, very high confidence; 70–90, confident; 50–70, low confidence; and <50, very low confidence. The resulting structural representation was used for illustration of the computationally predicted membrane orientation of NEMP-1 and was not interpreted as experimental determination of its transmembrane topology.

### Statistical analysis

All statistical analyses were performed in RStudio using R version 4.5.2. as noted in individual figure legends. All two sample Welch t-tests are unpaired and two-tailed. P values < 0.05 were considered statistically significant. For analyses of homolog pairing, synapsis, CHK-2 activity, and nuclear collapse across meiotic progression, three animals were analyzed per condition for each zone. Proportions were compared between conditions using Fisher’s exact test. Additional details of the statistical analyses are provided in the corresponding figure legends.

## Acknowledgements

We thank Abby Dernburg for reagents and computational resources, and Yumi Kim for pHIM/ZIMs and SYP-2 affinity-purified antibodies. We thank Makiah Brewer and Oluwatoyosi (Naomi) Adelola for assistance with the fertility and viability assay, and Ruijun Zhu for insightful discussions. We thank members of the Liu Lab for stimulating discussions and critically reading the manuscript. Some *C. elegans* strains used in this work were provided by the Caenorhabditis Genetics Center, which is funded by the NIH – Office of Research Infrastructure Programs (P40 OD010440).

## Funding

This work was supported by a startup fund from Lehigh University to CL, the Ralph E. Powe Junior Faculty Enhancement Award from Oak Ridge Associated Universities to CL, and a grant from the National Institutes of Health (R35GM160292) to CL.

## Author contributions

Conceptualization: CL; Methodology: RN, CL; Investigation: RN, CL; Visualization: RN, CL; Supervision: CL; Project administration: CL; Writing—original draft: RN, CL; Writing—review and editing: RN, CL; Funding acquisition: CL.

## Competing interests

Authors declare that they have no competing interests.

## Data and materials availability

All data needed to evaluate the conclusions in the paper are present in the paper and/or the Supplementary Materials. Strains generated in this study can be provided upon request.

## Supplementary Materials

Figure S1 and Figure S2; Tables S1 to S5; Video 1 and 2.

## Supplementary Materials

**Figure S1.**
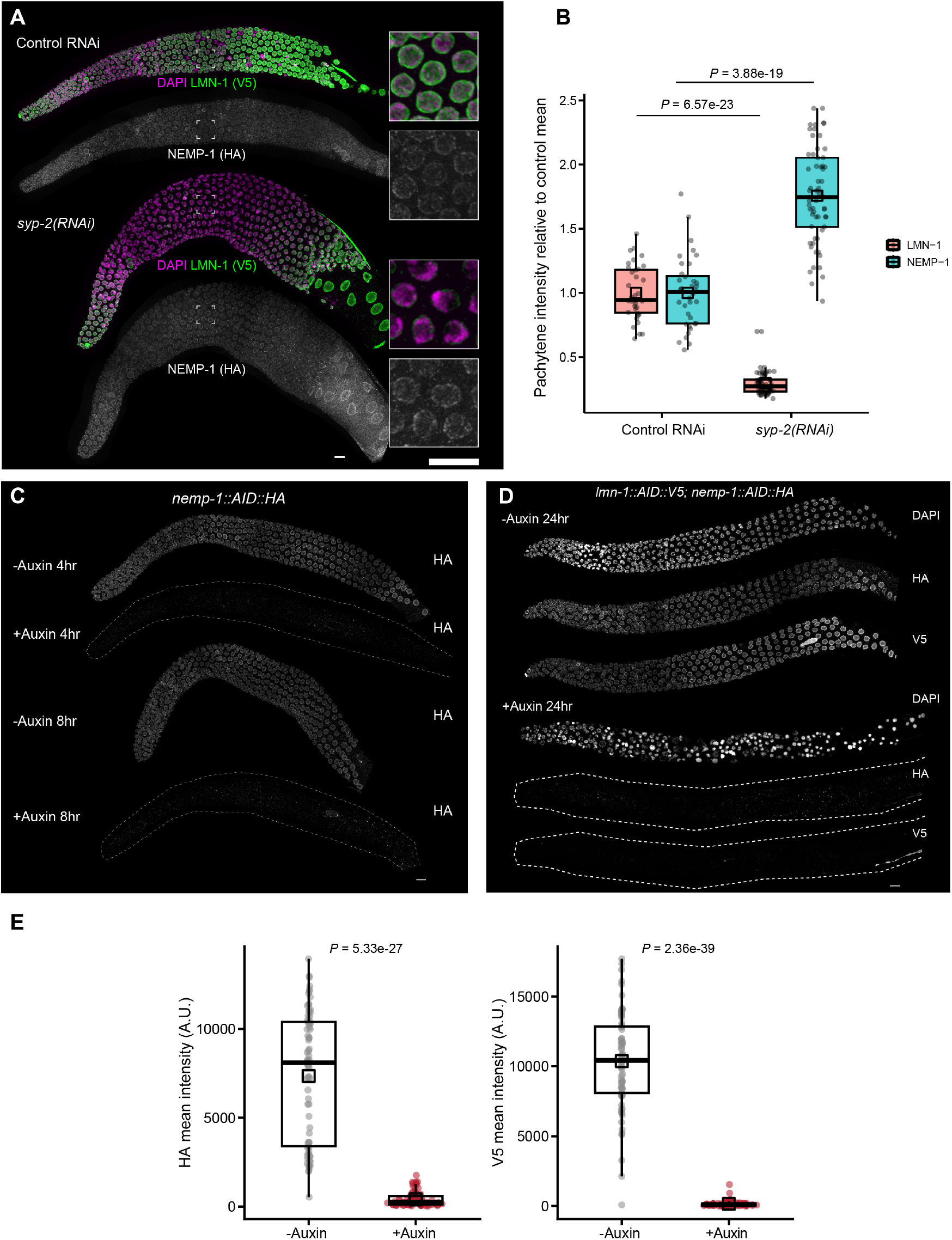
NEMP-1 localizes to the meiotic nuclear envelope and can be depleted using auxin-inducible degradation. (A) Immunofluorescence images showing NEMP-1 (HA staining) and LMN-1 (V5 staining) following syp-2(RNAi)-induced synapsis failure. DAPI in magenta, LMN-1 (V5) in green, and NEMP-1 (HA) in gray. Scale bars, 10 µm. (B) Quantification of LMN-1 and NEMP-1 intensities at the NE in mid-pachytene following control RNAi or syp-2(RNAi) treatment. LMN-1 and NEMP-1 fluorescence intensities were each normalized to the control group. Each point represents an individual nucleus. Two-tailed unpaired Welch’s t-test was used to compute P values. 36 nuclei were pooled from three animals for control RNAi, 61 nuclei were pooled from three animals for syp-2(RNAi). Exact P values are indicated in the figure. (C) Immunofluorescence images showing the effects on NEMP-1 (HA) staining following auxin-inducible degradation. Scale bar, 10 µm. (D) Immunofluorescence images showing the effects on the staining of NEMP-1 (HA) and LMN-1 (V5) following auxin-inducible degradation. Scale bar, 10 µm. (E) Quantification of HA and V5 intensities without or with auxin treatment for 24 h from young adult stage. 76 nuclei were pooled from 3 control animals. 80 nuclei were pooled from 3 auxin-treated animals. Additive projection images followed by background subtraction were used for measuring HA or V5 intensity with binary masks generated using the DAPI channel. Each point represents one nucleus, with Tukey boxplots overlaid and open squares indicating means. Unpaired Welch t-test was used to compute P values. Exact P values are indicated in the figure.

**Figure S2.**
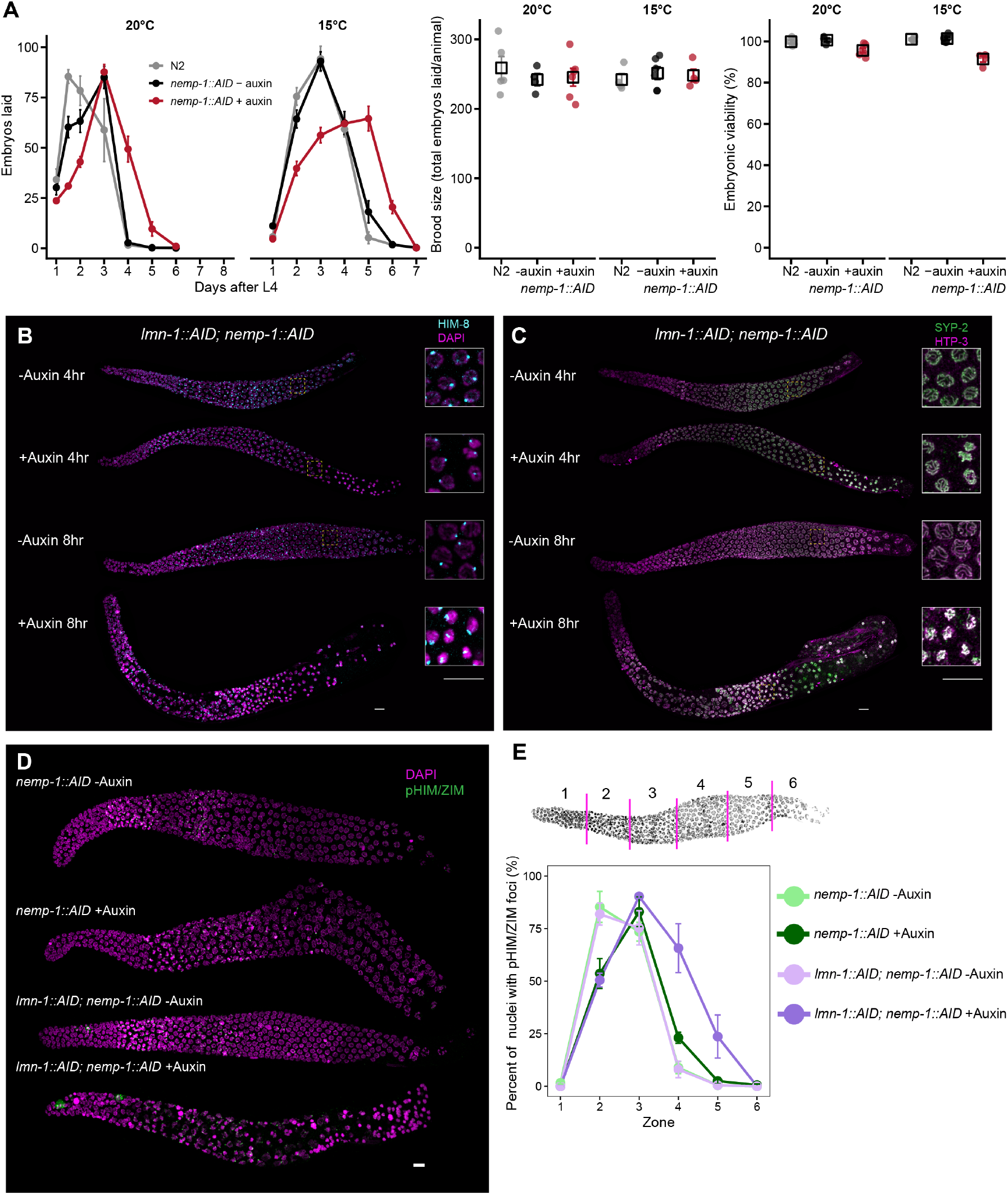
Effects of auxin-inducible degradation of NEMP-1 or LMN-1 in *C. elegans* germline. (A) Effects of NEMP-1 depletion on the timing of egg laying, brood size, and embryonic viability. Egg production was monitored following L1-onset auxin treatment of nemp-1::AID animals maintained at 20°C or 15°C. Left, number of eggs laid during each collection interval plotted according to days after the L4 stage. Points and error bars indicate mean ± SEM. Egg-laying profiles differed significantly between −auxin and +auxin animals at both 20°C and 15°C (condition × time interaction, repeated-measures ANOVA: 20°C, P = 3.97e−13; 15°C, P = 1.07e−12). Post hoc two-tailed Welch’s t-tests with Holm correction identified significant differences at day 4 at 20°C (P = 0.00469), and at days 2, 3, 5, and 6 at 15°C (P = 0.0127, 0.00207, 0.00474, and 0.0285, respectively). Middle, total egg production per animal. NEMP-1 depletion did not significantly alter total egg production at either 20°C (P = 0.833) or 15°C (P = 0.810). Right, embryonic viability, calculated for each animal as the total number of F1 progeny divided by the total number of eggs laid. NEMP-1 depletion significantly reduced embryonic viability at both 20°C (P = 0.00895) and 15°C (P = 0.00273). For middle and right panels, individual points represent animals, open squares indicate means, and error bars indicate SEM; comparisons were performed using two-tailed unpaired Welch’s t-tests. Detailed counts are provided in Table S1. (B and C) Immunofluorescence images showing the effects of auxin-inducible degradation of both NEMP-1 and LMN-1 on homolog pairing and synapsis. For (B), immunofluorescence images showing X-chromosome homolog pairing visualized by HIM-8 staining. HIM-8 in cyan and DAPI in magenta. For (C), immunofluorescence images showing chromosome synapsis visualized by SYP-2 and HTP-3 staining. SYP-2 in green and HTP-3 in magenta. Scale bars, 10 µm. (D) Immunofluorescence images showing phosphorylated HIM-8/ZIM proteins (pHIM/ZIM), a readout of CHK-2 activity. pHIM/ZIM in green and DAPI in magenta. Scale bar, 10 µm. (E) Quantification of the percent of nuclei with pHIM/ZIM foci as a function of meiotic progression. Distal gonad was divided into six equal-length zones, and the percent of nuclei with pHIM/ZIM foci in each zone was quantified. Three animals were analyzed per condition, and data are plotted as mean ± SEM. Nuclear counts were pooled across animals for pairwise comparisons of proportions. Comparisons are made for light green vs. dark green and light purple vs. dark purple, respectively: zone 2: P = 3.11e-15 and 3.01e-09; zone 3: P = 1.27e-02 and 4.90e-04; zone 4: P = 1.11e-05 and 2.55e-26; zone 5: P = 1.22e-01 and 2.48e-09; zone 6: P = 1.00 and 1.00.

## Supplementary Movies

**Movie S1.**
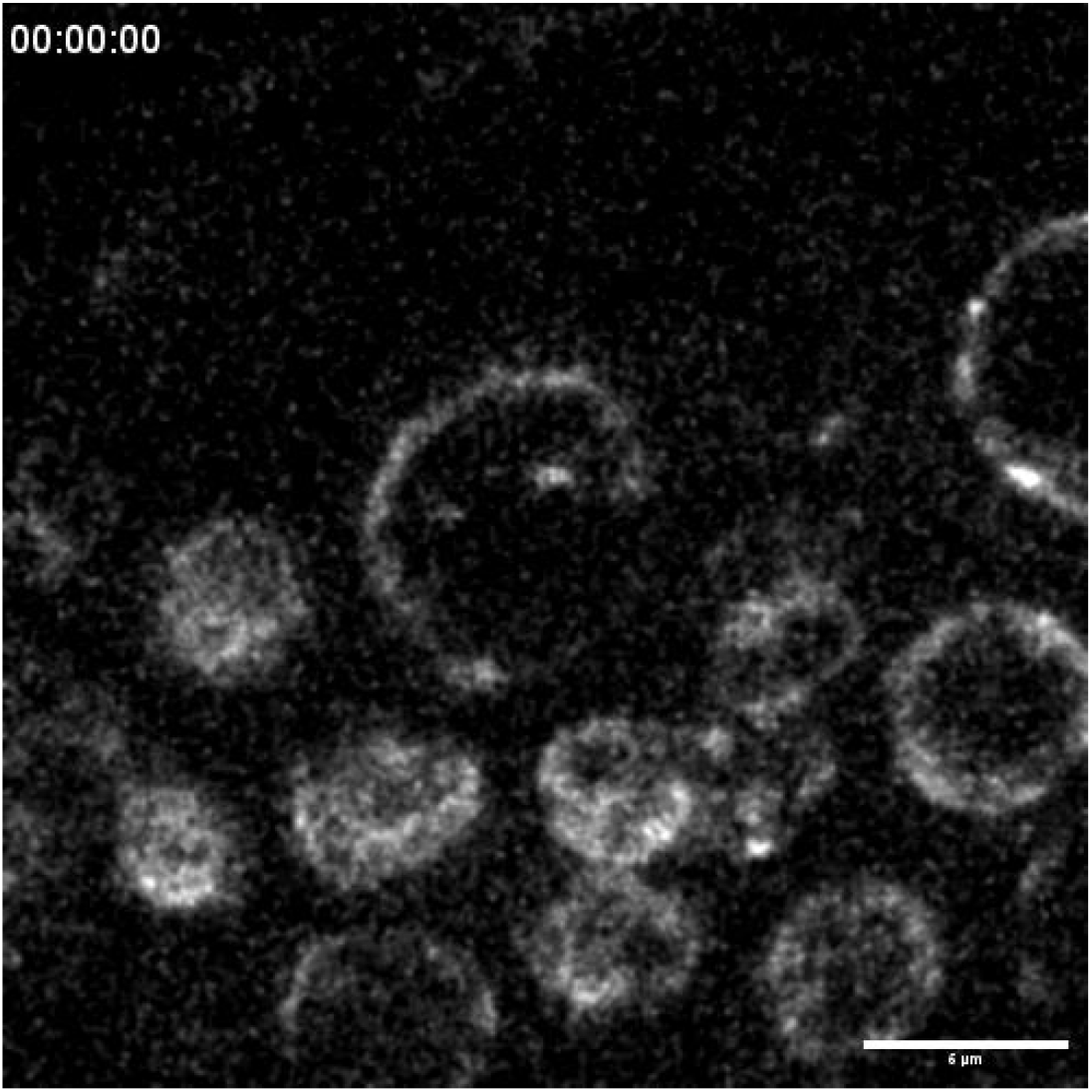
Simultaneous depletion of both NEMP-1 and LMN-1 results in acute nuclear collapse in early pachytene. Time-lapse recording of a representative nuclear collapse event in early pachytene following simultaneous depletion of NEMP-1 and LMN-1. 4D image stack of ZYG-12::GFP was used for generating maximum-intensity Z-projection at each time point. Time stamp is h:min:s. Scale bar, 5 µm.

**Movie S2.**
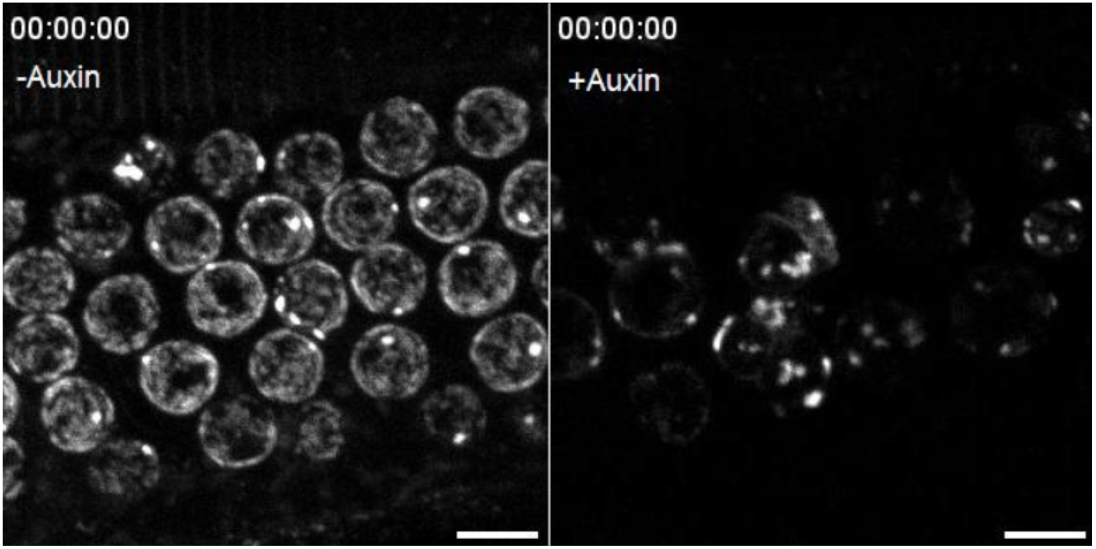
Depleting both NEMP-1 and LMN-1 enhances LINC dynamics at the NE in early pachytene. Time-lapse recording of the dynamics of LINC complex patches/clusters at the NE in early pachytene. A 3D image stack of ZYG-12::GFP was used for generating maximum-intensity Z-projection at each time point. Control group (-Auxin) is on the left and auxin treated group (+Auxin) is on the right. Time stamp is h:min:s. Scale bar, 5 µm.

**Table S1.** Quantification of brood size, embryonic viability, and male self-progeny of worm strains with alleles generated in this study. Sample size n indicates the number of broods examined.

| Strains | n | Condition | Embryos laid (Mean ±SD) | Embryonic viability (Mean ±SD) %* | Male progeny (Mean ±SD) % |
| --- | --- | --- | --- | --- | --- |
| N2 | 5 | - auxin, 20°C | 258.8 ± 37.1 | 99.8 ± 1.7 | 0.2 ± 0.2 |
| *ieSi64[gld-1p::TIR1::mRuby::gld-13'UTR, Cbr-unc-119(+)] II; unc-119(ed3), nemp-1(emj1[nemp-1::AID::HA]) III* | 4 | - auxin, 20°C | 241.8 ± 17.0 | 100.7 ± 1.7 | 0.3 ± 0.2 |
| *ieSi64[gld-1p::TIR1::mRuby::gld-13'UTR, Cbr-unc-119(+)] II; unc-119(ed3), nemp-1(emj1[nemp-1::AID::HA]) III* | 6 | + auxin (from L1), 20°C | 244.3 ± 30.3 | 95.6 ± 3.0 | 0.1 ± 0.2 |
| N2 | 4 | - auxin, 15°C | 241.8 ± 17.0 | 101.0 ± 0.9 | 0.3 ± 0.2 |
| *ieSi64[gld-1p::TIR1::mRuby::gld-13'UTR, Cbr-unc-119(+)] II; unc-119(ed3), nemp-1(emj1[nemp-1::AID::HA]) III* | 6 | - auxin, 15°C | 250.8 ± 21.8 | 101.3 ± 1.5 | 0.1 ± 0.2 |
| *ieSi64[gld-1p::TIR1::mRuby::gld-13'UTR, Cbr-unc-119(+)] II; unc-119(ed3), nemp-1(emj1[nemp-1::AID::HA]) III* | 4 | + auxin (from L1), 15°C | 247.8 ± 18.1 | 91.4 ± 3.0 | 0.1 ± 0.2 |

\* Embryonic viability of >100% reflects the fact that some embryos are overlooked when counting, but adult worms hatched from these embryos are easier to count accurately.

**Table S2.**
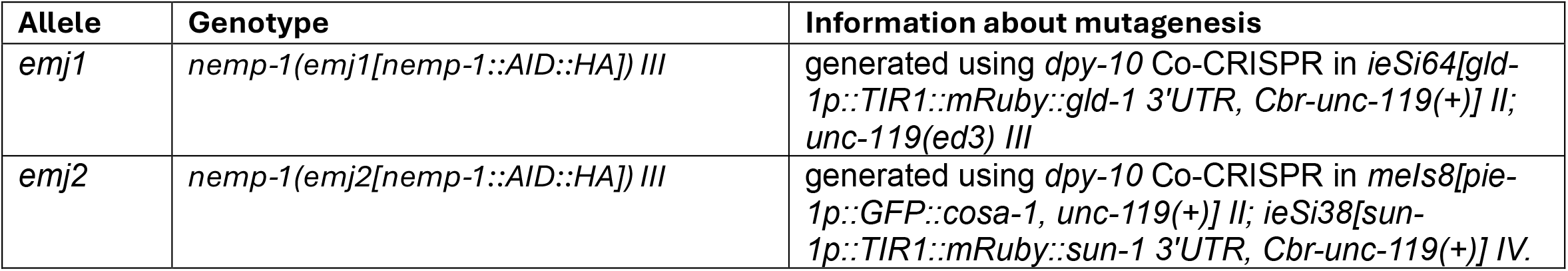
Alleles generated in this study.

| Allele | Genotype | Information about mutagenesis |
| --- | --- | --- |
| *emj1* | *nemp-1(emj1[nemp-1::AID::HA]) III* | generated using *dpy-10* Co-CRISPR in *ieSi64[gld-1p::TIR1::mRuby::gld-1 3'UTR, Cbr-unc-119(+)] II; unc-119(ed3) III* |
| *emj2* | *nemp-1(emj2[nemp-1::AID::HA]) III* | generated using *dpy-10* Co-CRISPR in *mels8[pie-1p::GFP::cosa-1, unc-119(+)] II; ieSi38[sun-1p::TIR1::mRuby::sun-1 3'UTR, Cbr-unc-119(+)] IV.* |

**Table S3.** Genotypes of worm strains generated and used in this study.

| Strain | Source | Identifier |
| --- | --- | --- |
| *C. elegans*: N2 Bristol, wild isolate | Caenorhabditis Genetics Center | N2 |
| *C. elegans*: *ieSi64*[*gld-1p::TIR1::mRuby::gld-1 3'UTR*, *Cbr-unc-119(+)*] II; *unc-119(ed3)* III | Zhang et al., 2015; Caenorhabditis Genetics Center | CA1352 |
| *C. elegans*: *mels8*[*pie-1p::GFP::cosa-1*, *unc-119(+)*] II; *ieSi38*[*sun-1p::TIR1::mRuby::sun-1 3'UTR*, *Cbr-unc-119(+)*] IV | Zhang et al., 2018 | CA1364 |
| *C. elegans*: *lmn-1*(*ie137*[*lmn-1::AID::V5*]) I; *unc-119 (ed3)* III; *ieSi38* [*sun-1p::TIR1::mRuby::sun-1 3'UTR*, *cb-unc-119(+)*] IV | Liu et al., 2023 | CEL6 (CA1532) |
| *C. elegans*: *unc-119 (ed3)* III; *ieSi38* [*sun-1p::TIR1::mRuby::sun-1 3'UTR*, *cb-unc-119(+)*] IV; *sun-1*(*ie139*[*sun-1::AID::V5*]) V | Liu et al., 2023 | CEL38 (CA1564) |
| *C. elegans*: *lmn-1*(*ie137*[*lmn-1::AID::V5*]) I; *ieSi64*[*gld-1p::TIR1::mRuby::gld-1 3'UTR*, *Cbr-unc-119(+)*] II; *unc-119(ed3)* III; *ieSi19* (*mRuby::SYP-3*),*ojls9* [*zyg-12(all)::GFP + unc-119(+)*] IV | Liu et al., 2023 | CEL48 (CA1574) |
| *C. elegans*: *ieSi64*[*gld-1p::TIR1::mRuby::gld-13'UTR*, *Cbr-unc-119(+)*] II; *unc-119(ed3)*, *nemp-1*(*emj1*[*nemp-1::AID::HA*]) III | This paper | CEL272 |
| *C. elegans*: *lmn-1*(*ie137*[*lmn-1::AID::V5*]) I; *nemp-1*(*emj2*[*nemp-1::AID::HA*]) III; *ieSi38*[*sun-1p::TIR1::mRuby::sun-1 3'UTR*, *Cbr-unc-119(+)*] IV | This paper | CEL278 |
| *C. elegans*: *lmn-1*(*ie137*[*lmn-1::AID::V5*]) I; *nemp-1*(*emj2*[*nemp-1::AID::HA*]) III; *ieSi38*[*sun-1p::TIR1::mRuby::sun-1 3'UTR*, *Cbr-unc-119(+)*] IV; *sun-1*(*ie139*[*sun-1::AID::V5*]) V | This paper | CEL281 |
| *C. elegans*: *lmn-1*(*ie137*[*lmn-1::AID::V5*]) I; *ieSi64*[*gld-1p::TIR1::mRuby::gld-13'UTR*, *Cbr-unc-119(+)*] II; *unc-119(ed3)*, *nemp-1*(*emj2*[*nemp-1::AID::HA*]) III; *ieSi19* (*mRuby::SYP-3*),*ojls9* [*zyg-12(all)::GFP + unc-119(+)*] IV | This paper | CEL286 |
| *C. elegans*: *ieSi64*[*gld-1p::TIR1::mRuby::gld-13'UTR*, *Cbr-unc-119(+)*] II; *unc-119(ed3)*, *nemp-1*(*emj1*[*nemp-1::AID::HA*]) III; *ieSi19* (*mRuby::SYP-3*),*ojls9* [*zyg-12(all)::GFP + unc-119(+)*] IV | This paper | CEL318 |

**Table S4.**
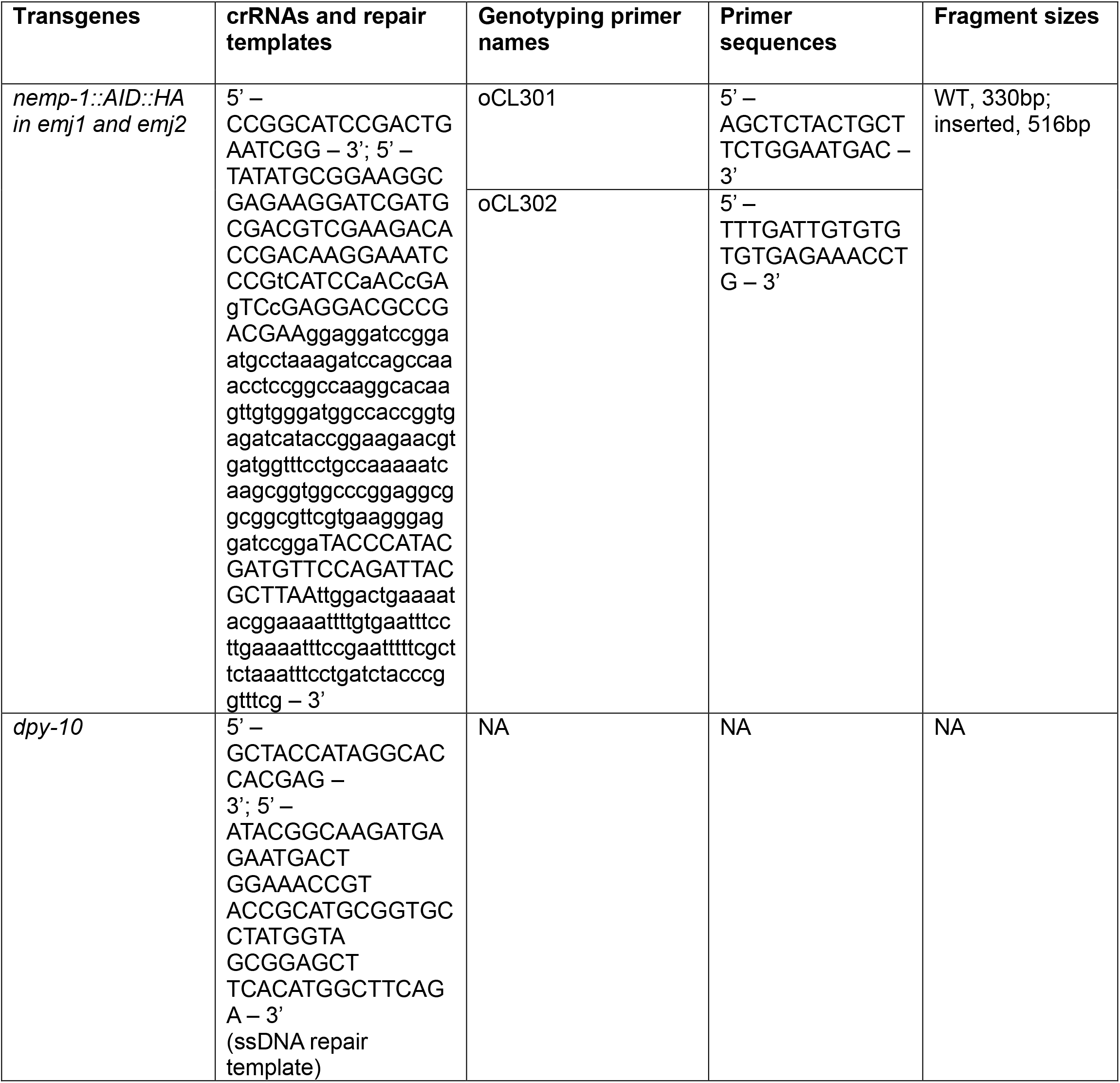
Sequences of crRNAs, repair templates, and DNA primers used to genotype edited progeny.

**Table S5.** RNAi clones used in this study. Both RNAi constructs were confirmed using Nanopore sequencing of whole-plasmids (Plasmidsaurus).

| Target gene | GenePairs name | Plate | Well |
| --- | --- | --- | --- |
| *syp-2* | C24G6.1 | 138 | B6 |
| *dnc-1* | ZK593.5 | 111 | B12 |

## Notes

### Competing Interest Statement

The authors have declared no competing interest.

## References

1. D. Zickler, N. Kleckner, Meiosis: Dances between homologs, Annual Review of Genetics 57 (Volume 57, 2023) (2023) 1–63. doi:10.1146/annurev-genet-061323-044915. URL https://www.annualreviews.org/content/journals/10.1146/annurev-genet-061323-044915

2. N. Bhalla, A. F. Dernburg, Prelude to a division, Annual review of cell and developmental biology 24 (2008) 397–424. doi:10.1146/annurev.cellbio.23.090506.123245. URL http://www.ncbi.nlm.nih.gov/pmc/articles/PMC4435778/

3. I. R. Adams, O. R. Davies, Meiotic chromosome structure, the synaptonemal complex, and infertility, Annual Review of Genomics and Human Genetics 24 (Volume 24, 2023) (2023) 35–61. doi:10.1146/annurev-genom-110122-090239. URL https://www.annualreviews.org/content/journals/10.1146/annurev-genom-110122-090239

4. B. I. Cesar, Y. Kim, Structure and function of the synaptonemal complex, Journal of Cell Biology 225 (5) (2026) e202511222. doi:10.1083/jcb.202511222. URL https://doi.org/10.1083/jcb.202511222

5. C. M. Lake, R. S. Hawley, Synaptonemal complex, Current Biology 31 (5) (2021) R225– R227. doi:10.1016/j.cub.2021.01.015. URL https://doi.org/10.1016/j.cub.2021.01.015

6. K. S. McKim, K. Peters, A. M. Rose, Two types of sites required for meiotic chromosomepairing in caenorhabditis elegans, Genetics 134 (3) (1993) 749–768. doi:10.1093/genetics/134.3.749. URL https://doi.org/10.1093/genetics/134.3.749

7. O. Rog, A. F. Dernburg, Chromosome pairing and synapsis during caenorhab-ditis elegans meiosis, Current Opinion in Cell Biology 25 (3) (2013) 349–356. doi:10.1016/j.ceb.2013.03.003. URL http://www.sciencedirect.com/science/article/pii/S095506741300046X

8. A. M. Villeneuve, A cis-acting locus that promotes crossing over between x chromo-somes in caenorhabditis elegans, Genetics 136 (3) (1994) 887–902. doi:10.1093/genetics/136.3.887. URL https://doi.org/10.1093/genetics/136.3.887

9. A. F. Dernburg, Pushing the (nuclear) envelope into meiosis, Genome Biology 14 (3) (2013) 1–3. doi:10.1186/gb-2013-14-3-110. URL http://dx.doi.org/10.1186/gb-2013-14-3-11010.

10. (2022). doi:10.3390/genes13050901.

11. R. Nenstiel, C. Liu, Forceful Beginnings: Mechanotransduction and Mechanosensationat the Meiotic Cell Nucleus During Gamete Development, Springer Nature Switzerland, Cham, 2026, pp. 221–247. doi:10.1007/978-3-032-19509-8_10. URL https://doi.org/10.1007/978-3-032-19509-8_10

12. B. Cai, M. Tiscareno-Andrade, Y. Luo, S. Lefranc, F. Cao, A. Chambon, X. Yuan, M. Peuch, Y. Zhang, A. Hurel, J. Guérin, N. Vrielynck, C. Mézard, P. Andrey, L. Cromer, C. Yang, M. Grelon, Identification of the cytoplasmic motor–linc complex involved in rapid chromosome movements during meiotic prophase in arabidopsis thaliana, Nature Plants 11 (8) (2025) 1608–1627. doi:10.1038/s41477-025-02043-4. URL https://doi.org/10.1038/s41477-025-02043-4

13. Y. Chikashige, C. Tsutsumi, M. Yamane, K. Okamasa, T. Haraguchi, Y. Hiraoka, Meiotic proteins bqt1 and bqt2 tether telomeres to form the bouquet arrangement of chromo-somes, Cell 125 (1) (2006) 59–69. doi:10.1016/j.cell.2006.01.048. URL https://doi.org/10.1016/j.cell.2006.01.048

14. N. Christophorou, T. Rubin, I. Bonnet, T. Piolot, M. Arnaud, J.-R. Huynh, Microtubule-driven nuclear rotations promote meiotic chromosome dynamics, Nature Cell Biology 17 (11) (2015) 1388–1400. doi:10.1038/ncb3249. URL https://doi.org/10.1038/ncb3249

15. M. N. Conrad, C.-Y. Lee, G. Chao, M. Shinohara, H. Kosaka, A. Shinohara, J. A. Conchello, M. E. Dresser, Rapid telomere movement in meiotic prophase is promoted by <em>ndj1</em>, <em>mps3</em>, and <em>csm4</em> and is modulated by re-combination, Cell 133 (7) (2008) 1175–1187. doi:10.1016/j.cell.2008.04.047. URL https://doi.org/10.1016/j.cell.2008.04.047

16. L. Cromer, M. Tiscareno-Andrade, S. Lefranc, A. Chambon, A. Hurel, M. Brogniez, J. Guérin, I. Le Masson, G. Adam, D. Charif, P. Andrey, M. Grelon, Rapid meiotic prophase chromosome movements in arabidopsis thaliana are linked to essential re-organization at the nuclear envelope, Nature Communications 15 (1) (2024) 5964. doi:10.1038/s41467-024-50169-4. URL https://doi.org/10.1038/s41467-024-50169-4

17. D.-Q. Ding, A. Yamamoto, T. Haraguchi, Y. Hiraoka, Dynamics of homologous chromo-some pairing during meiotic prophase in fission yeast, Developmental Cell 6 (3) (2004) 329–341. doi:10.1016/S1534-5807(04)00059-0. URL https://doi.org/10.1016/S1534-5807(04)00059-0

18. X. Ding, R. Xu, J. Yu, T. Xu, Y. Zhuang, M. Han, Sun1 is required for telomere at-tachment to nuclear envelope and gametogenesis in mice, Developmental Cell 12 (6) (2007) 863–872. doi:10.1016/j.devcel.2007.03.018. URL https://doi.org/10.1016/j.devcel.2007.03.018

19. H. J. Kim, C. Liu, L. Zhang, A. F. Dernburg, Mjl-1 is a nuclear envelope protein required for homologous chromosome pairing and regulation of synapsis during meiosis in c. elegans, Science Advances 9 (6) (2023) eadd1453. doi:10.1126/sciadv.add1453. URL https://doi.org/10.1126/sciadv.add1453

20. C.-Y. Lee, H. Horn, C. Stewart, B. Burke, E. Bolcun-Filas, J. Schimenti, M. Dresser, R. Pezza, Mechanism and regulation of rapid telomere prophase movements in mouse meiotic chromosomes, Cell Reports 11 (4) (2015) 551–563. doi:10.1016/j.celrep.2015.03.045. URL https://www.sciencedirect.com/science/article/pii/S2211124715003241

21. Q. Meng, B. Shao, D. Zhao, X. Fu, J. Wang, H. Li, Q. Zhou, T. Gao, Loss of sun1 function in spermatocytes disrupts the attachment of telomeres to the nuclear envelope and contributes to non-obstructive azoospermia in humans, Human Genetics 142 (4) (2023) 531–541. doi:10.1007/s00439-022-02515-z. URL https://doi.org/10.1007/s00439-022-02515-z

22. A. Morimoto, H. Shibuya, X. Zhu, J. Kim, K.-i. Ishiguro, M. Han, Y. Watanabe, A conserved kash domain protein associates with telomeres, sun1, and dynactin dur-ing mammalian meiosis, Journal of Cell Biology 198 (2) (2012) 165–172. doi:10.1083/jcb.201204085. URL https://doi.org/10.1083/jcb.201204085

23. A. Mytlis, V. Kumar, T. Qiu, R. Deis, N. Hart, K. Levy, M. Masek, A. Shawahny, A. Ah-mad, H. Eitan, F. Nather, S. Adar-Levor, R. Y. Birnbaum, N. Elia, R. Bachmann-Gagescu, S. Roy, Y. M. Elkouby, Control of meiotic chromosomal bouquet and germ cell morphogenesis by the zygotene cilium, Science 376 (6599) (2022) eabh3104. doi:10.1126/science.abh3104. URL https://doi.org/10.1126/science.abh3104

24. A. Penkner, L. Tang, M. Novatchkova, M. Ladurner, A. Fridkin, Y. Gruenbaum, D. Schweizer, J. Loidl, V. Jantsch, The nuclear envelope protein matefin/sun-1 is required for homologous pairing in c. elegans meiosis, Developmental Cell 12 (6) (2007) 873–885. doi:10.1016/j.devcel.2007.05.004. URL http://www.sciencedirect.com/science/article/pii/S1534580707001992

25. A. Sato, B. Isaac, C. M. Phillips, R. Rillo, P. M. Carlton, D. J. Wynne, R. A. Kasad, A. F. Dernburg, Cytoskeletal forces span the nuclear envelope to coordinate meiotic chromosome pairing and synapsis, Cell 139 (5) (2009) 907–919. doi:10.1016/j.cell.2009.10.039. URL http://www.sciencedirect.com/science/article/pii/S0092867409013658

26. A. Viera, M. Alsheimer, R. Gómez, I. Berenguer, S. Ortega, C. E. Symonds, D. San-tamaría, R. Benavente, J. A. Suja, Cdk2 regulates nuclear envelope protein dynamics and telomere attachment in mouse meiotic prophase, Journal of Cell Science 128 (1) (2015) 88–99. doi:10.1242/jcs.154922. URL https://doi.org/10.1242/jcs.154922

27. J. J. Wanat, K. P. Kim, R. Koszul, S. Zanders, B. Weiner, N. Kleckner, E. Alani, Csm4, in collaboration with ndj1, mediates telomere-led chromosome dynamics and recombination during yeast meiosis, PLOS Genetics 4 (9) (2008) e1000188. doi: 10.1371/journal.pgen.1000188. URL https://doi.org/10.1371/journal.pgen.1000188

28. C. M. Phillips, C. Wong, N. Bhalla, P. M. Carlton, P. Weiser, P. M. Meneely, F. Dernburg, Him-8 binds to the x chromosome pairing center and mediates chromosome-specific meiotic synapsis, Cell 123 (6) (2005) 1051–1063. doi:10.1016/j.cell.2005.09.035. URL http://www.sciencedirect.com/science/article/pii/S009286740501041X

29. D. J. Wynne, O. Rog, P. M. Carlton, A. F. Dernburg, Dynein-dependent processive chromosome motions promote homologous pairing in c. elegans meiosis, The Journal of Cell Biology 196 (1) (2012) 47–64. URL http://jcb.rupress.org/content/196/1/47.abstract

30. D. Y. Lui, M. P. Colaiácovo, Meiotic Development in Caenorhabditis elegans, Springer New York, New York, NY, 2013, pp. 133–170. doi:10.1007/978-1-4614-4015-4_6. URL https://doi.org/10.1007/978-1-4614-4015-4_6

31. C. M. Phillips, K. L. McDonald, A. F. Dernburg, Cytological Analysis of Meiosis in Caenorhabditis elegans, Humana Press, Totowa, NJ, 2009, pp. 171–195. doi: 10.1007/978-1-60761-103-5_11. URL https://doi.org/10.1007/978-1-60761-103-5_11

32. C. M. Phillips, A. F. Dernburg, A family of zinc-finger proteins is required for chromosome-specific pairing and synapsis during meiosis in c. elegans, Develop-mental Cell 11 (6) (2006) 817–829. doi:10.1016/j.devcel.2006.09.020. URL http://www.sciencedirect.com/science/article/pii/S1534580706004503

33. N. Harper, R. Rillo, S. Jover-Gil, Z. Assaf, N. Bhalla, A. Dernburg, Pairing centers recruit a polo-like kinase to orchestrate meiotic chromosome dynamics in c. elegans, Developmental Cell 21 (5) (2011) 934–947. URL http://linkinghub.elsevier.com/retrieve/pii/S1534580711003996

34. O. Rog, A. Dernburg, Direct visualization reveals kinetics of meiotic chromosome synapsis, Cell Reports 10 (10) (2015) 1639–1645. doi:10.1016/j.celrep.2015.02.032. URL http://www.sciencedirect.com/science/article/pii/S2211124715001783

35. G. Huelgas-Morales, M. Sanders, G. Mekonnen, T. Tsukamoto, D. Greenstein, De-creased mechanotransduction prevents nuclear collapse in a lt;emgt;caenorhabditis eleganslt;/emgt; laminopathy, Proceedings of the National Academy of Sciences 117 (49) (2020) 31301. doi:10.1073/pnas.2015050117. URL http://www.pnas.org/content/117/49/31301.abstract

36. C. Liu, R. Rex, Z. Lung, J. S. Wang, F. Wu, H. J. Kim, L. Zhang, L. L. Sohn, A. F. Dernburg, A cooperative network at the nuclear envelope counteracts linc-mediated forces during oogenesis in c. elegans, Science Advances 9 (28) (2023) eabn5709. doi:10.1126/sciadv.abn5709. URL https://doi.org/10.1126/sciadv.abn5709

37. C. Liu, A. F. Dernburg, Chemically induced proximity reveals a piezo-dependent meiotic checkpoint at the oocyte nuclear envelope, Science 386 (6724) (2024) eadm7969. doi: 10.1126/science.adm7969. URL https://doi.org/10.1126/science.adm7969

38. J. Link, D. Paouneskou, M. Velkova, A. Daryabeigi, T. Laos, S. Labella, C. Barroso, S. Pacheco Piñol, A. Montoya, H. Kramer, A. Woglar, A. Baudrimont, S. M. Markert, C. Stigloher, E. Martinez-Perez, A. Dammermann, M. Alsheimer, M. Zetka, V. Jantsch, Transient and partial nuclear lamina disruption promotes chromosome movement in early meiotic prophase, Developmental Cell 45 (2) (2018) 212–225.e7. doi:10.1016/j.devcel.2018.03.018. URL https://doi.org/10.1016/j.devcel.2018.03.018

39. N. Bhalla, A. F. Dernburg, A conserved checkpoint monitors meiotic chromosome synapsis in caenorhabditis elegans, Science 310 (5754) (2005) 1683–1686. URL http://science.sciencemag.org/content/310/5754/1683.abstract

40. A. Gartner, S. Milstein, S. Ahmed, J. Hodgkin, M. O. Hengartner, A conserved check-point pathway mediates dna damagex2013;induced apoptosis and cell cycle arrest in <em>c. elegans</em>, Molecular Cell 5 (3) (2000) 435–443. doi:10.1016/S1097-2765(00)80438-4. URL https://doi.org/10.1016/S1097-2765(00)80438-4

41. T. L. Gumienny, E. Lambie, E. Hartwieg, H. R. Horvitz, M. O. Hengartner, Genetic control of programmed cell death in the caenorhabditis elegans hermaphrodite germline, Development 126 (5) (1999) 1011–1022. doi:10.1242/dev.126.5.1011. URL https://doi.org/10.1242/dev.126.5.1011

42. Y. Tsatskis, R. Rosenfeld, J. D. Pearson, C. Boswell, Y. Qu, K. Kim, L. Fabian, A. Mo-hammad, X. Wang, M. I. Robson, K. Krchma, J. Wu, J. Gonçalves, D. Hodzic, S. Wu, D. Potter, L. Pelletier, W. H. Dunham, A.-C. Gingras, Y. Sun, J. Meng, D. Godt, T. Schedl, B. Ciruna, K. Choi, J. R. B. Perry, R. Bremner, E. C. Schirmer, J. A. Brill, A. Jurisicova, H. McNeill, The nemp family supports metazoan fertility and nuclear envelope stiffness, Science Advances 6 (35) (2020) eabb4591. doi:10.1126/sciadv.abb4591. URL http://advances.sciencemag.org/content/6/35/eabb4591.abstract

43. Y. Kim, N. Kostow, A. Dernburg, The chromosome axis mediates feedback control of chk-2 to ensure crossover formation in c. elegans, Developmental Cell 35 (2) (2015) 247–261. doi:10.1016/j.devcel.2015.09.021. URL http://www.sciencedirect.com/science/article/pii/S153458071500622X

44. S. Rosu, K. A. Zawadzki, E. L. Stamper, D. E. Libuda, A. L. Reese, A. F. Dernburg, M. Villeneuve, The c. elegans dsb-2 protein reveals a regulatory network that con-trols competence for meiotic dsb formation and promotes crossover assurance, PLOS Genetics 9 (8) (2013) e1003674. doi:10.1371/journal.pgen.1003674. URL https://doi.org/10.1371/journal.pgen.1003674

45. E. L. Stamper, S. E. Rodenbusch, S. Rosu, J. Ahringer, A. M. Villeneuve, A. F. Dern-burg, Identification of dsb-1, a protein required for initiation of meiotic recombination in caenorhabditis elegans, illuminates a crossover assurance checkpoint, PLOS Genetics 9 (8) (2013) e1003679. doi:10.1371/journal.pgen.1003679. URL https://doi.org/10.1371/journal.pgen.1003679

46. L. Zhang, S. Köhler, R. Rillo-Bohn, A. F. Dernburg, A compartmentalized signaling network mediates crossover control in meiosis, eLife 7 (2018) e30789. doi:10.7554/eLife.30789. URL https://doi.org/10.7554/eLife.30789

47. L. Zhang, J. D. Ward, Z. Cheng, A. F. Dernburg, The auxin-inducible degradation (aid) system enables versatile conditional protein depletion in c. elegans, Development 142 (24) (2015) 4374–4384. URL http://dev.biologists.org/content/142/24/4374.abstract

48. A. Baudrimont, A. Penkner, A. Woglar, T. Machacek, C. Wegrostek, J. Gloggnitzer, A. Fridkin, F. Klein, Y. Gruenbaum, P. Pasierbek, V. Jantsch, Leptotene/zygotene chro-mosome movement via the sun/kash protein bridge in caenorhabditis elegans, PLOS Genetics 6 (11) (2010) e1001219. doi:10.1371/journal.pgen.1001219. URL https://doi.org/10.1371/journal.pgen.1001219

49. A. M. Penkner, A. Fridkin, J. Gloggnitzer, A. Baudrimont, T. Machacek, A. Woglar, E. Csaszar, P. Pasierbek, G. Ammerer, Y. Gruenbaum, V. Jantsch, Meiotic chromosome homology search involves modifications of the nuclear envelope protein matefin/sun-1, Cell 139 (5) (2009) 920–933. doi:10.1016/j.cell.2009.10.045. URL http://dx.doi.org/10.1016/j.cell.2009.10.045

50. T. Bohr, G. Ashley, E. Eggleston, K. Firestone, N. Bhalla, Synaptonemal complex components are required for meiotic checkpoint function in caenorhabditis elegans, Genetics 204 (3) (2016) 987–997. doi:10.1534/genetics.116.191494. URL https://pubmed.ncbi.nlm.nih.gov/27605049 https://www.ncbi.nlm.nih.gov/pmc/articles/PMC5105873/

51. M. Castellano-Pozo, S. Pacheco, G. Sioutas, A. L. Jaso-Tamame, M. H. Dore, M. M. Karimi, E. Martinez-Perez, Surveillance of cohesin-supported chromosome structure controls meiotic progression, Nature Communications 11 (1) (2020) 4345. doi:10.1038/s41467-020-18219-9. URL https://doi.org/10.1038/s41467-020-18219-9

52. A. Woglar, A. Daryabeigi, A. Adamo, C. Habacher, T. Machacek, A. La Volpe, V. Jantsch, Matefin/sun-1 phosphorylation is part of a surveillance mechanism to co-ordinate chromosome synapsis and recombination with meiotic progression and chro-mosome movement, PLOS Genetics 9 (3) (2013) e1003335. doi:10.1371/journal.pgen.1003335. URL https://doi.org/10.1371/journal.pgen.1003335

53. D. J. Barbosa, J. Duro, B. Prevo, D. K. Cheerambathur, A. X. Carvalho, R. Gassmann, Dynactin binding to tyrosinated microtubules promotes centrosome centration in c. ele-gans by enhancing dynein-mediated organelle transport, PLOS Genetics 13 (7) (2017) e1006941. doi:10.1371/journal.pgen.1006941. URL https://doi.org/10.1371/journal.pgen.1006941

54. A. R. Skop, J. G. White, The dynactin complex is required for cleavage plane spec-ification in early caenorhabditis elegans embryos, Current Biology 8 (20) (1998) 1110–1117. doi:10.1016/S0960-9822(98)70465-8. URL https://www.sciencedirect.com/science/article/pii/S0960982298704658

55. M. Terasawa, M. Toya, F. Motegi, M. Mana, K. Nakamura, A. Sugimoto, Caenorhabditis elegans ortholog of the p24/p22 subunit, dnc-3, is essential for the formation of the dynactin complex by bridging dnc-1/p150glued and dnc-2/dynamitin, Genes to Cells 15 (11) (2010) 1145–1157. doi:10.1111/j.1365-2443.2010.01451.x. URL https://doi.org/10.1111/j.1365-2443.2010.01451.x

56. R. B. Vallee, R. J. McKenney, K. M. Ori-McKenney, Multiple modes of cytoplasmic dynein regulation, Nat Cell Biol 14 (3) (2012) 224–230. URL http://dx.doi.org/10.1038/ncb2420

57. H. Zhang, A. R. Skop, J. G. White, Src and wnt signaling regulate dynactin accumulation to the p2-ems cell border in c. elegans embryos, Journal of Cell Science 121 (2) (2008) 155–161. doi:10.1242/jcs.015966. URL https://doi.org/10.1242/jcs.015966

58. M. P. Colaiácovo, A. J. MacQueen, E. Martinez-Perez, K. McDonald, A. Adamo, La Volpe, A. M. Villeneuve, Synaptonemal complex assembly in <em>c. ele-gans</em> is dispensable for loading strand-exchange proteins but critical for proper completion of recombination, Developmental Cell 5 (3) (2003) 463–474. doi:10.1016/S1534-5807(03)00232-6. URL https://doi.org/10.1016/S1534-5807(03)00232-6

59. A. J. Deshong, A. L. Ye, P. Lamelza, N. Bhalla, A quality control mechanism coordinates meiotic prophase events to promote crossover assurance, PLOS Genetics 10 (4) (2014) e1004291. doi:10.1371/journal.pgen.1004291. URL https://doi.org/10.1371/journal.pgen.1004291

60. A. J. MacQueen, M. P. Colaiácovo, K. McDonald, A. M. Villeneuve, Synapsis-dependent and -independent mechanisms stabilize homolog pairing during meiotic prophase in c. elegans, Genes Development 16 (18) (2002) 2428–2442. URL http://genesdev.cshlp.org/content/16/18/2428.abstract

61. A. J. MacQueen, C. M. Phillips, N. Bhalla, P. Weiser, A. M. Villeneuve, F. Dernburg, Chromosome sites play dual roles to establish homologous synapsis during meiosis in c. elegans, Cell 123 (6) (2005) 1037–1050. doi:10.1016/j.cell.2005.09.034. URL http://www.sciencedirect.com/science/article/pii/S0092867405010408

62. S. Smolikov, K. Schild-Prüfert, M. P. Colaiácovo, A yeast two-hybrid screen for syp-3 interactors identifies syp-4, a component required for synaptonemal complex assembly and chiasma formation in caenorhabditis elegans meiosis, PLOS Genetics 5 (10) (2009) e1000669. doi:10.1371/journal.pgen.1000669. URL https://doi.org/10.1371/journal.pgen.1000669

63. B. A. Hakim, Y. Tsatskis, L. Zhang, E. Choi, Y. Zhang, D. Hodzic, Q. Wu, M. Zhang, M. Pashaei, K. Ha, J. Rusch, J. A. Brill, M. A. Brieño-Enríquez, A. Jurisicova, H. McNeill, Loss of nemp1 disrupts female meiosis and activates a conserved atm-chk2 checkpoint, Nature Communications 17 (1) (2026) 9100. doi:10.1038/s41467-026-75874-0. URL https://doi.org/10.1038/s41467-026-75874-0

64. J. Link, V. Jantsch, Meiotic chromosomes in motion: a perspective from mus musculus and caenorhabditis elegans, Chromosoma 128 (3) (2019) 317–330. doi:10.1007/s00412-019-00698-5. URL https://doi.org/10.1007/s00412-019-00698-5

65. H. Shibuya, Telomeres, the nuclear lamina, and membrane remodeling: Orchestrating meiotic chromosome movements, Journal of Cell Biology 224 (5) (2025) e202412135. doi:10.1083/jcb.202412135. URL https://doi.org/10.1083/jcb.202412135

66. W. Xie, M. Gowder, D. Bazzano, M. DeSantis, S. S. Hammoud, Rewiring for movements in meiotic prophase: regulators, roles, and evolutionary pathways, Current Opinion in Genetics Development 93 (2025) 102366. doi:10.1016/j.gde.2025.102366. URL https://www.sciencedirect.com/science/article/pii/S0959437X25000589

67. L. Zhao, J. Gao, Chromosome movements in meiotic prophase: regulatory mechanisms and biological roles, Frontiers in Cell and Developmental Biology Volume 14 - 2026 (2026). URL https://www.frontiersin.org/journals/cell-and-developmental-biology/articles/10.3389/fcell.2026.1911284

68. S. Brenner, The genetics of caenorhabditis elegans, Genetics 77 (1) (1974) 71–94. URL https://pubmed.ncbi.nlm.nih.gov/4366476 https://www.ncbi.nlm.nih.gov/pmc/articles/PMC1213120/

69. T. Davies, H. X. Kim, N. Romano Spica, B. J. Lesea-Pringle, J. Dumont, M. Shirasu-Hiza, J. C. Canman, Cell-intrinsic and -extrinsic mechanisms promote cell-type-specific cytokinetic diversity, eLife 7 (2018) e36204. doi:10.7554/eLife.36204. URL https://doi.org/10.7554/eLife.36204

70. L. Timmons, D. L. Court, A. Fire, Ingestion of bacterially expressed dsrnas can produce specific and potent genetic interference in caenorhabditis elegans, Gene 263 (1) (2001) 103–112. doi:10.1016/S0378-1119(00)00579-5. URL https://www.sciencedirect.com/science/article/pii/S0378111900005795

71. R. S. Kamath, J. Ahringer, Genome-wide rnai screening in caenorhabditis elegans, Methods 30 (4) (2003) 313–321. doi:10.1016/S1046-2023(03)00050-1. URL https://www.sciencedirect.com/science/article/pii/S1046202303000501

72. R. S. Kamath, A. G. Fraser, Y. Dong, G. Poulin, R. Durbin, M. Gotta, A. Kanapin, N. Le Bot, S. Moreno, M. Sohrmann, D. P. Welchman, P. Zipperlen, J. Ahringer, Systematic functional analysis of the caenorhabditis elegans genome using rnai, Nature 421 (6920) (2003) 231–237. doi:10.1038/nature01278. URL https://doi.org/10.1038/nature01278

73. M. E. HURLock, I. Č avka, L. E. Kursel, J. Haversat, M. Wooten, Z. Nizami, R. Turniansky, P. Hoess, J. Ries, J. G. Gall, O. Rog, S. Köhler, Y. Kim, Identification of novel synaptonemal complex components in c. elegans, Journal of Cell Biology 219 (5) (2020) e201910043. doi:10.1083/jcb.201910043. URL https://doi.org/10.1083/jcb.201910043

74. C. Liu, Y. Mao, Diaphanous formin mdia2 regulates cenp-a levels at centromeres, The Journal of Cell Biology 213 (4) (2016) 415–424. URL http://jcb.rupress.org/content/213/4/415.abstract

75. A. L. Lomize, S. C. Todd, I. D. Pogozheva, Spatial arrangement of proteins in planar and curved membranes by ppm 3.0, Protein Science 31 (1) (2022) 209–220. doi:10.1002/pro.4219. URL https://doi.org/10.1002/pro.4219

